# Single capsid mutations modulating phage adsorption, persistence, and plaque morphology shape evolutionary trajectories in ΦX174

**DOI:** 10.1101/2025.09.09.674889

**Authors:** Manuela Reuter, Michael Sieber, Octavio Reyes-Matte, Christina Vasileiou, Christopher Böhmker, Jordan Romeyer Dherbey, Frederic Bertels, Javier Lopez-Garrido

## Abstract

The evolutionary success of lytic bacteriophages depends on key life-history traits, including adsorption, lysis time, burst size, and persistence in the environment. However, how these traits evolve to allow adaptation to different environments remains poorly understood. Here, we explored this question in ΦX174, combining experimental evolution and mathematical modelling. By investigating how serial transfer conditions shape evolutionary outcomes in liquid culture, we found that the time between transfers imposes divergent selection on adsorption and context-dependent directional selection on persistence. Longer transfer intervals, which allow multiple infection cycles until host depletion, favoured fast-adsorbing, highly persistent mutants that could rapidly initiate infections and remained viable in the absence of the host. In contrast, shorter transfer intervals selected for slower adsorption without substantially altering persistence. Mathematical modelling of phage population dynamics predicted that adsorption evolution during short transfers reflects a trade-off between two opposing selective forces within each transfer: an early phase in which susceptible hosts are abundant and adsorption is productive, favouring fast adsorption, and a later phase in which most hosts are already infected and adsorption primarily removes phage particles via attachment to already infected cells, favouring slower adsorption. A single-point mutation in the major capsid protein was sufficient to drive these changes in adsorption. In the case of fast-adsorbing mutants, this mutation was positively pleiotropic and also enhanced environmental persistence. Our findings show how simple changes in propagation conditions can steer phage phenotypes, providing insights relevant to evolutionary biology and phage therapy.

## Introduction

The success of lytic bacteriophages relies on four central life-history traits, each of which may evolve under different selective pressures to maximise fitness [1–6]. These traits are: (I) adsorption constant, which determines the ease with which a phage irreversibly attaches to a host cell, (II) latent period, or time required to complete the infection cycle inside the host, (III) burst size, defined as the number of new phage particles produced per infected host cell, and (IV) decay rate, representing the loss of infectious phage particles in the absence of susceptible host cells [7–12].

Interest in these traits has grown due to the need to predict phage virulence for medical and biocontrol applications [13–16]. However, such predictions remain challenging because the fitness effects of individual traits are often highly context-dependent [17–19]. For example, high adsorption rates are typically considered beneficial in well-mixed liquid environments, where they facilitate rapid infection initiation [20, 21]. However, in spatially-structured environments such as biofilms, these same traits can hinder phage spread to uninfected neighbouring cells by promoting phage attachment to already infected cells at the centre of infection [17, 22]. Understanding how life-history traits influence phage fitness across environments is therefore essential for predicting and potentially optimising phage performance.

The model coliphage ΦX174 provides a powerful platform for dissecting how the relationship between trait and environment impacts fitness [23–27]. ΦX174 is a strictly lytic, tailless phage of the *Microviridae* family, best known as the first DNA-based biological entity whose entire genome was sequenced [28]. Its 5386 bp circular single-stranded DNA genome encodes only 11 genes, most of which are necessary for phage DNA replication, morphogenesis, and host lysis [28–31]. Only four genes (F, G, H, and J) encode for proteins that form the mature virion [32, 33]. The icosahedral capsid is made of 60 F, 60 J and 12 H protein monomers, decorated at each of its 12 vertices by 5 G spike monomers [32, 34]. G likely mediates host recognition and subsequent attachment to host lipopolysaccharide (LPS), after which it dissociates, allowing F to bind irreversibly to the host cell surface [35–37]. Irreversible binding triggers a conformational change that results in the injection of phage DNA into the host cell through the pilot protein H [37, 38].

Here, we have explored how life-history traits influence phage fitness across different environments by evolving ΦX174 under different serial transfer regimes. Through a combination of life-history trait quantification and mathematical modelling, we have identified individual mutations that alter adsorption constant and persistence, leading to fitness effects that are strongly dependent on the environment. By disentangling these context-dependent fitness effects, our results improve our understanding of phage ecology and evolution and also provide insights for rational trait optimisation in medical and biocontrol applications.

## Materials and Methods

### Strains and culture conditions

The ancestral ΦX174 and its host *E. coli* C were provided by H.A. Wichman (Department of Biological Sciences, University of Idaho). All evolved phage strains were derived from the same ΦX174 genotype (GenBank: AF176034). Cultures were grown in Lysogeny Broth (LB, Miller) supplemented with 10 mM MgCl₂ and 5 mM CaCl₂ and incubated at 37°C, shaking at 250 rpm (orbital shaking diameter 2.5 cm). Mid-exponential *E. coli* C cultures (∼1-2*10^8^ colony-forming units [CFU] per ml) were prepared from overnight cultures and used for all transfer experiments and life history quantification assays.

### Phage lysate preparation and titration

Phage lysates were generated as described in Romeyer Dherbey *et al.* [39] by co-culturing exponentially growing *E. coli* C with ΦX174 for 3 hours in a final volume of 5 ml. After 3 hours, the remaining bacteria were removed with 10-12 drops of chloroform and subsequent centrifugation at 5000 rpm for 10 minutes at 4°C. Phage-containing supernatants were collected and stored as lysate at 4°C or at −80°C as glycerol stocks for long-term storage.

Phage titers were quantified by plaque assays in which plaque-forming units (PFUs) were assessed by plating 100 µl of serially diluted phage lysate with 100 µl of *E. coli* C stationary-phase culture pre-mixed in 4 ml semi-solid agar (SSA, 0.5% agar, supplemented with 10 mM MgCl_2_ and 5 mM CaCl_2_). The mixture was overlayed on a plate of LB-agar (1.5%) and incubated at 37°C for 18 hours. Phage lysates were diluted in phage buffer juice (PBJ: 2.03% Tris-HCl, 0.61% MgCl_2_).

### Serial transfer regimes

We implemented two transfer regimes, the 3-h regime and the 30-min regime, which are described in detail below. For both transfer regimes, five independent selection lines were initiated by infecting mid-exponential phage-sensitive *E. coli* C cultures (∼1-2*10^8^ CFU/ml) with ancestral ΦX174 extracted from independent plaques. Each transfer was performed in a final volume of 5 ml of supplemented LB medium (see “Strains and culture conditions”). All transfer cultures were incubated at 37°C, shaking at 250 rpm (orbital shaking diameter 2.5 cm). The 3-h and 30-min regimes are described in detail below. Additional details on the transfer regimes can be found in **Fig. S1**.

### 3-h transfer regime

The 3-h regime was chosen based on a previous finding in our lab, which showed that small plaques begin to emerge after approximately seven 3-hour transfers of ancestral ΦX174 [39]. This regime was initiated by infecting a 5-ml culture of exponentially growing *E. coli* C (∼1-2*10^8^ CFU/ml) at an MOI of ∼0.1, followed by a 3-h incubation at 37°C with shaking at 250 rpm. After the incubation period, phages were extracted with chloroform. The resulting phage lysate was titrated and used to inoculate the next transfer on the following day at MOI_input_ of ∼0.1 on a fresh, mid-exponential host culture (∼1-2*10^8^ CFU/ml) (**Fig. 1**; **Fig. S1A**). We repeated this cycle every day and performed titration assays after every transfer to quantify titers and screen for changes in plaque morphology. Serial transfers were continued until heritable small-plaque mutants emerged. When detected, these small plaques were isolated and replated to confirm the phenotype.

**Fig. 1:**
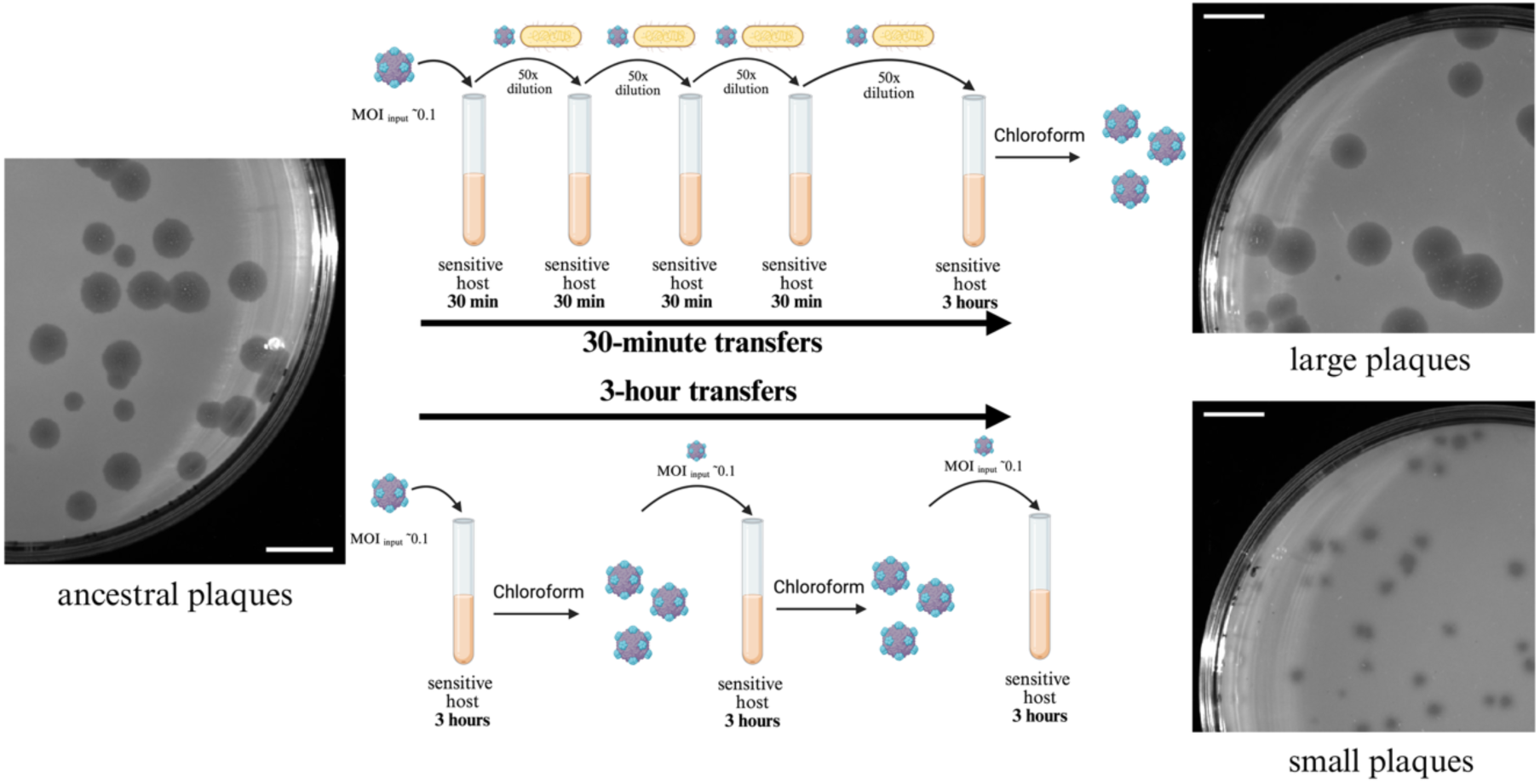
30-min and 3-h transfer regimes select for large- and small-plaque-forming phages, respectively. Phage ΦX174 was propagated under 30-min (top) or 3-h (bottom) regimes, leading to the evolution of large- and small-plaque-forming phages, respectively. Ancestral plaques are shown on the left, and evolved large (top) and small (bottom) plaques on the right. Scale bars, 1 cm. All plaque images have the same magnification. Diagrams of the 30-min and 3-h regimes are in the middle. The approximate input multiplicity of infection (MOI_input_) and incubation times are the same for each transfer. For illustration purposes, only four transfers are shown, but the experiment involved more transfers. For the 30-min regime, 20 transfers (fifteen 30-min transfers, interrupted by a 3-h transfer after every fourth transfer) were done before large-plaque mutants were observed in plaque assays. In the 3-h regime, transfers were done until heritable small plaques emerged, which occurred after four to seven transfers. See methods and **Fig. S1** for details about the individual transfer protocols. Figure created in BioRender. Reuter, M. (2025) https://BioRender.com/q39e4in.

### 30-min transfer regime

The 30-min regime was a modification of the 3-h regime in which 3-h transfers were interspersed by blocks of four 30-min transfers (**Fig. 1**; **Fig. S1B**). The first 30-min transfer of each block was initiated by infecting a 5-ml culture of exponentially growing *E. coli* C (∼1-2*10^8^ CFU/ml) at an MOI of ∼0.1. The culture was incubated for 30 min at 37°C with shaking at 250 rpm, after which 2% of the phage-bacteria co-culture was transferred to 5 ml of a fresh, mid-exponential *E. coli* C culture. This transfer was repeated after each 30-min incubation for a total of four consecutive 30-min transfers. At the end of the fourth 30-min transfer, 2% of the phage-bacteria co-culture was transferred to 5 ml of a fresh, mid-exponential *E. coli* C culture and incubated for 3h at 37°C with shaking at 250 rpm. After this incubation, phages were extracted with chloroform. The lysate was titrated and used to seed the next 30-min transfer block on the following day. Plaques larger than the ancestral ones emerged in all five lineages (see results). Phage lysate of each transfer day was stored for Sanger sequencing.

### Sanger sequencing of evolved phages

Whole-genome Sanger sequencing was conducted for individual phage isolates from both transfer regimes according to Biggs *et al.* [40]. DNA was extracted from high-titer lysates (QIAprep Spin Miniprep Kit® Qiagen, Hilden, Germany) and amplified via PCR in two genome halves. PCR products were purified (QIAprep PCR Purification Kit® Qiagen, Hilden, Germany) and sequenced at the Max Planck Institute for Evolutionary Biology. Primers and settings used for amplification and sequencing are in **Tables S1, S2, and S3**. Sequences were aligned against the reference ΦX174 genome using Geneious Prime 2023.0.1 (https://www.genious.com). Identified amino acid positions in which single-point mutations were located were numbered according to the residue number, which differs from the codon number by minus one because the first amino acid (methionine) of the F protein is cleaved [32, 41]. Raw Sanger Sequencing files are deposited in Zenodo (see Zenodo doi: 10.5281/zenodo.17076220).

### Adsorption assays

Adsorption constants were determined following a modified version of the Hyman and Abedon assay [42]. Mid-exponential, phage-sensitive *E. coli* C cultures were co-cultured with phages at MOI_input_ ∼0.05 at 37°C in a shaking water bath at 200 rpm. Phages were sampled 2, 4, and 6 minutes after adsorption initiation and diluted 10-fold into non-supplemented LB. Samples were cooled on ice to avoid temperature-dependent decay effects, treated with chloroform to stop infection, and quantified. Adsorption constants were calculated by fitting linear models to log-transformed phage titers and dividing slope estimates by bacterial density. Differences between genotypes were tested using a linear mixed model (replicate as a random effect) and post hoc Tukey tests for multiple comparisons (**Supplementary Methods**).

### Phage persistence assays

Persistence was measured over a 5-hour co-culturing period of ΦX174 and an exponentially-growing *E. coli* C culture infected with phages at MOI_input_ ∼0.1. Phage samples were taken every 30 minutes, extracted with chloroform, and phage titers were determined by PFU counting. Decay rates were estimated by fitting exponential decay models to the titer decline period after peak productivity. Model fitting procedures are detailed in the **Supplementary Methods**.

### Simulations of phage evolution

The following system of delay differential equations (DDEs) describes the dynamics of the bacterial host 𝐵 (cells per ml), the infected bacterial population 𝐵_*I*_(cells per ml), and the three phage types 𝑃_*A*_ (ancestral, particles per ml), 𝑃_*L*_ (large-plaque type evolved in vitro in the 30-min regime, particles per ml), and 𝑃_*S*_ (small-plaque type evolved in vitro in the 3-h regime, particles per ml). In addition, 𝐵_𝜏_ = 𝐵(𝑡 − 𝜏) denotes the bacterial density 𝜏 minutes in the past, reflecting the delay introduced by the lysis time. We use a similar notation for the phage densities, e.g. 𝑃_*A*, 𝜏_ denotes the density of the ancestral phage 𝜏 minutes in the past. For 𝑡 < 𝜏, we set the corresponding densities to zero. Annotations of small and large plaque types were based on the in-vitro phenotypes specified in the “Serial Transfer Section” to simplify labelling. However, plaque morphology was not included in the in-silico modelling.

Bacterial growth throughout the transfer period was found not to play a significant role in the simulations, so it is neglected in the model for simplicity. All parameter values for phage life traits and their biological meaning are given in **Table 1**. Beyond the values given in Table 1, we tested the robustness of the in-silico models for the 30-min and 3-h regimes to variation in central phage life history traits (i.e. lysis time and burst size) (**Fig. S2**).

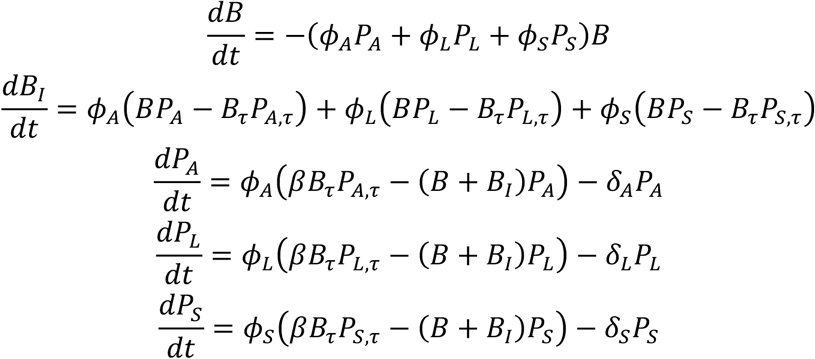

**Table 1:** Parameters and corresponding parameter values used in the mathematical model.

| Parameter | Value |
| --- | --- |
| Adsorption constant, ancestor $\Phi_A$ | $4 \cdot 10^{-9} \text{ (ml}^{-1} \text{ min}^{-1}\text{)}$ |
| Adsorption constant, large-plaque type evolved in vitro in the 30-min regime $\Phi_L$ | $2 \cdot 10^{-9} \text{ (ml}^{-1} \text{ min}^{-1}\text{)}$ |
| Adsorption constant, small-plaque type evolved in vitro in the 3-h regime $\Phi_S$ | $6 \cdot 10^{-9} \text{ (ml}^{-1} \text{ min}^{-1}\text{)}$ |
| Burst size, all types $\beta$ | 200 (per infected host cell) |
| Lysis time, all types $\tau$ | 17 min |
| Decay rate, ancestor $\delta$ | $1.7 \cdot 10^{-2} \text{ (min}^{-1}\text{)}$ |
| Decay rate, ancestor and large-plaque type $\delta_L$ | $1.4 \cdot 10^{-2} \text{ (min}^{-1}\text{)}$ |
| Decay rate, small-plaque type $\delta_S$ | $6 \cdot 10^{-3} \text{ (min}^{-1}\text{)}$ |

The DDE system was solved numerically in Python using the *scipy*-based *ddeInt* solver. After 30 minutes or 3 hours of simulated time, depending on the transfer regime, the system was reset to the initial density of fresh bacteria 𝐵 (10^8^ CFU). In the 3-h regime, each transfer started with 10^7^ PFU from the previous transfer. In the 30-min regime, the first transfer was also initiated with 10^7^ PFU, and subsequent transfers with 2% of the final densities of infected bacteria 𝐵_!_. For the competition simulations, phages with different life history traits were introduced at different initial densities, as indicated in the corresponding figure legends. The model makes the following simplifying assumptions. First, it assumes a fixed lysis time of 17 min, according to our experimental calculations (**Fig. S3**). Second, it assumes that infected host cells disappear upon lysis and do not contribute to phage attachment as debris. Third, the model assumes a constant phage decay rate. The code to reproduce the figures is available at https://github.com/OReyesMatte/Reuter_2025/tree/main/Simulations.

### Image analysis of lysis plaques

Lysis plaque morphology was quantified using a custom image analysis pipeline. Plaques were imaged under standardised conditions (ChemiDoc: Bio-Rad, California, USA; 0.1s exposure time) and processed using Ilastik [43] for pixel classification based on intensity, texture, and edge features. Post-processing steps included erosion, whole-filling, and watershed transformation using SciPy [44] and scikit-image [45]. Each plaque was assigned an ID and analysed for area (cm²), distance to periphery (via distance transform), and grayscale intensity (overlaying mask on original image) (details see **Supplementary Methods**).

### Data analysis and visualisation

Statistical analysis of adsorption constants, model fitting for adsorption and decay, as well as all data visualisations, were performed in R (version 4.2.1) [46]. Details of the packages and scripts are in https://github.com/OReyesMatte/Reuter_2025/tree/main/Data_analysis and **Table S4**. The protein structure, resolved by McKenna *et al.* [32], shown in **Fig. 4A**, was taken from RCSB PDB (reference 2BPA: https://www.wwpdb.org/pdb?id=pdb_00002bpa), and the mutation visualisation was done in UCSF Chimera (version 1.18). All cartoons were created using BioRender® as indicated in the corresponding figure legends.

## Results

### Differences in serial transfer regimes drive the evolution of contrasting lysis plaque phenotypes

In the context of a previous experimental evolution study [39], we observed that small, turbid plaques frequently emerged when ΦX174 was serially transferred after 3-hour co-culturing periods with host cells in liquid, shaking conditions. To confirm this observation, we evolved multiple independent selection lines of the ancestral ΦX174 under the same selection regime (hereafter, 3-h regime) and plated the evolving phage populations onto lawns of phage-sensitive host cells at every transfer (**Fig. 1; Fig. S1A**). Small, turbid plaques appeared in all lines after four to seven transfers (**Fig. S4A**). In parallel, we tested a modified regime in which the 3-h incubation periods were interspersed by four consecutive 30-min transfers, each initiated by transferring 2% of the phage-bacteria mixture into fresh, sensitive host cells (hereafter, referred to as the 30-min regime). In contrast to the 3-h transfer regime, no small plaques were detected under the 30-min regime, even after 50 transfers (**Fig. 1, Fig. S1B**). However, we began to observe plaques larger than those formed by the ancestral phage after approximately 20 transfers (**Fig. S4B**).

### Large- and small-plaque phenotypes are caused by different mutations in the major capsid (F) protein

Since differences in plaque morphology often reflect underlying changes in life history traits [2, 17, 22, 47–49], we hypothesised that our experimental regimes were selecting for different life-history trait changes in the 3-h and 30-min regimes. To identify the genetic modifications leading to the small- and large-plaque phenotypes, we isolated five phages of each type, one per selection line, and sequenced their genomes. Each evolved isolate differed from the ancestor by only one or two single-point mutations (**Fig. 2**).

**Fig. 2:**
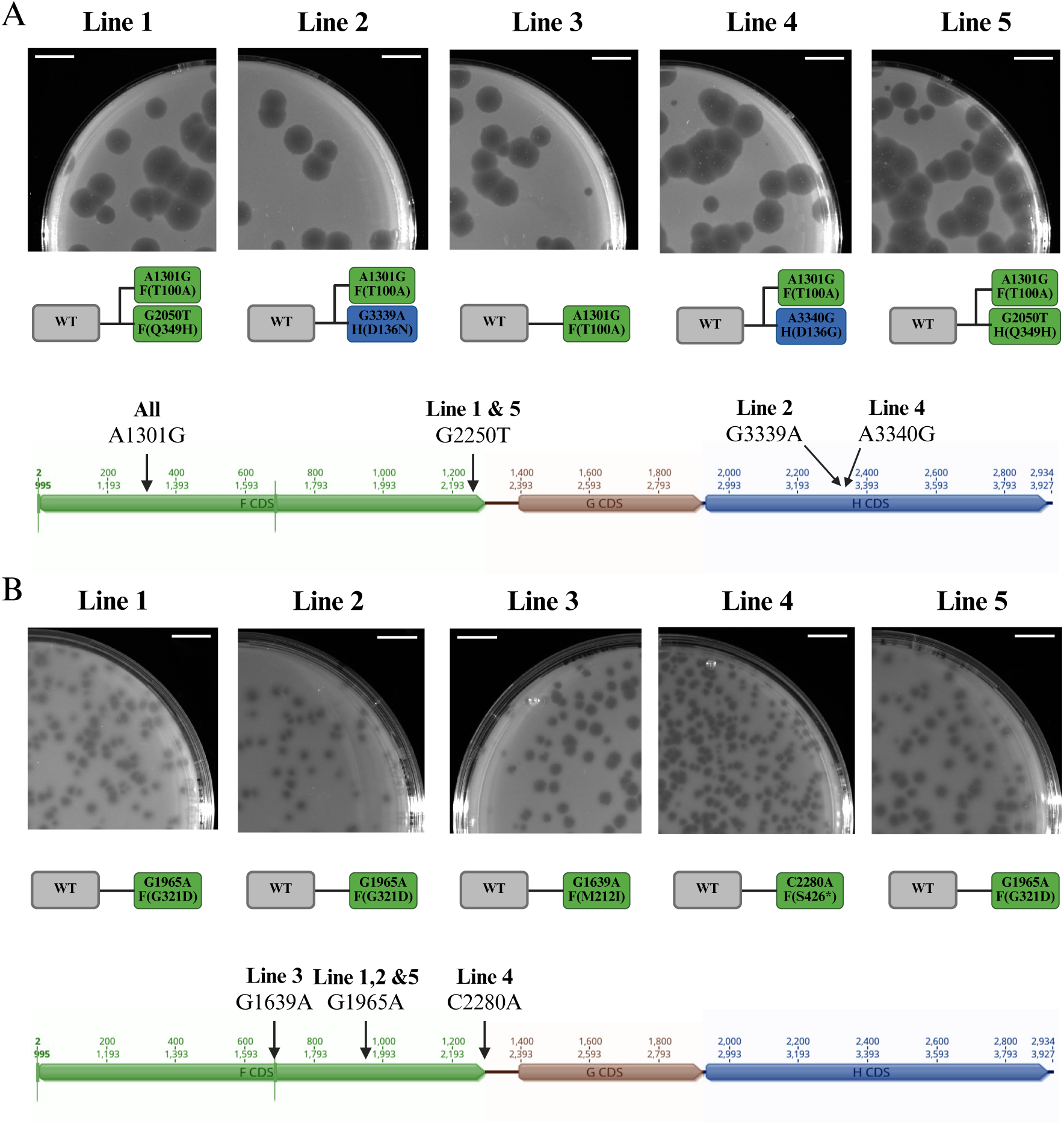
Mutations linked to large- and small-plaque phenotypes occur in the major capsid protein. (A) Large-plaque phenotypes (top) and corresponding genotypes (bottom) of mutants isolated from independent lines of the 30-min regime. All the mutants carried an A-to-G transition at position 1301 (A1301G) in the phage genome, resulting in a threonine-to-alanine substitution at position 101 of the F protein [F(T100A)]. Four isolates carried an additional mutation, either in the F (green) or the H (blue) gene. The ancestor is shown in grey (WT). (B) Small-plaque phenotypes (top) and genotypes (bottom) of mutants isolated from independent lines of the 3-h regime. All mutants differed from the ancestor by different mutation in the F gene (green), resulting in different amino acid changes: G321D in the clones from lines 1, 2, and 5; M212I in the clone from line 3; and the formation of a premature stop codon at the very last codon of the F gene (S426*) in the clone from line 4. The nucleotide change in the phage genome is shown at the top and the corresponding amino acid change at the bottom. The ancestor is shown in grey (WT). Scale bars, 1 cm. Genome illustrations were generated using Geneious Prime 2023.0.1. Figure created in BioRender. Reuter, M. (2025) https://BioRender.com/dck2411.

All five large-plaque mutants isolated from the 30-min regime lines carried an A-to-G transition at position 1301 in the phage genome, resulting in a threonine-to-alanine substitution at residue 100 of the F gene [A1301G/F(T100A); we use this notation throughout the manuscript: original nucleotide, nucleotide position, new nucleotide/protein (original amino acid, residue position, new amino acid)] (**Fig. 2A**). The mutant isolated from line 3 carried only this substitution. The other four isolates carried one additional mutation in either the F or the H protein (**Fig. 2A**) acquired after the F(T100A) mutation. Phenotypically, however, large-plaque-formers resembled each other, suggesting that the T100A mutation in the F protein is sufficient for the manifestation of the large-plaque phenotype.

None of the mutations observed in large-plaque mutants were found in the small-plaque-forming phages isolated from the 3-h regime lines. However, all five small-plaque phages also differed from the ancestor by a single-point mutation in the F gene (**Fig. 2B**), each producing a distinct small-plaque phenotype (**Fig. 2B; Fig. S5**). The most drastic phenotype was caused by the G1965A/F(G321D) mutation, resulting in the formation of extremely small and turbid plaques. Phages carrying the C2280A/F(S426*) mutation, which introduces an early stop codon and truncates the major capsid protein by the last amino acid, produced slightly larger and clearer plaques (**Fig. S5**). The G1639A/F(M212I) mutation led to a heterogeneous phenotype, with both small and large plaques, and was excluded from further analysis.

Altogether, the sequencing results indicate that different modifications of the major capsid protein are responsible for the contrasting plaque phenotypes emerging in our selection regimes.

### Plaque size negatively correlates with the adsorption constant

The major capsid protein is critical for the irreversible adsorption of ΦX174 to the host cells [37], suggesting the large- and small-plaque-forming phages may have evolved different adsorption constants compared to the ancestral ΦX174. To test this hypothesis, we measured the adsorption constant of the ancestral phage, a large-plaque-forming mutant carrying the single F(T100A) mutation evolved in the 30-min regime, and the two small-plaque mutants, F(G321D) and F(S426*), isolated from the 3-h regime. We focused on mutants with single substitutions to ensure that each mutation could be directly linked to specific changes in life history traits.

The adsorption constants of the evolved phages differed significantly from that of the ancestral ΦX174 and correlated inversely with plaque size (**Fig. 3**). The ancestral mean adsorption constant was 4.15×10^-9^ ml^-1^ min^-1^ (**Table S5**), in close agreement with previous estimations [50, 51]. The adsorption constants of the small-plaque-forming mutants evolved in the 3-h regime were slightly but significantly higher than those of the ancestral. Conversely, the large-plaque mutant evolved in the 30-min regime had an absorption rate constant approximately five-fold lower than the ancestral (**Table S5**).

**Fig. 3:**
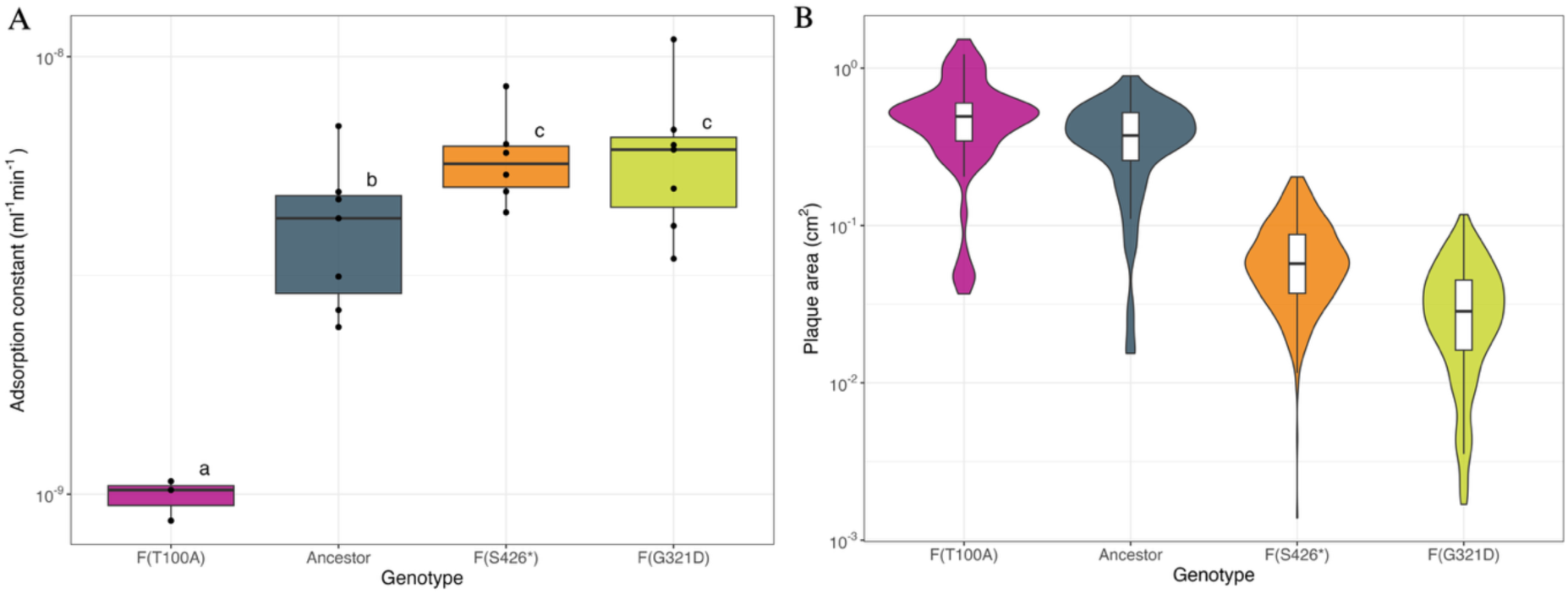
The adsorption constant is inversely correlated with plaque size. (A) Adsorption constant “k” of the ancestor (grey), the large-plaque mutant F(T100A) (purple) evolved from the 30-min regime and the two small-plaque mutants F(G321D) (orange) and F(S426*) isolated from the 3-h regime (green). At least three independent experiments were performed for each genotype. Each black dot represents the adsorption constant of an individual experiment, the horizontal lines indicate the median, and the boxes span the middle 50% of the data. The whiskers span the lower and upper 25% of the data. Different letters on top of the boxes indicate significant adsorption constant differences between genotypes (significance level p<0.05). (B) Lysis plaque area in cm^2^ (y-axis) measured in vitro for the ancestor (grey, n=76), and the evolved phages F(T100A) (purple, n=60, 30-min regime), F(G321D) (green, n=303, 3-h regime) and F(S426*) (orange, n=431, 3-h regime). Violin plots show distribution shape; boxplots display medians (black lines), interquartile range (boxes), and upper/lower 25% (whiskers). See **Methods** and **Supplementary Methods** for details of adsorption and plaque phenotype analysis.

The relationship between adsorption constant and plaque size was not linear. The large-plaque mutant with a five-fold reduction in adsorption constant produced plaques that were 1.5-fold larger than those of the ancestor, whereas the small-plaque mutants with less than two-fold increase in adsorption constant formed plaques about ten times smaller (**Table S5**). These results are in line with previous predictions of a non-linear relationship between adsorption constant and plaque size [49].

These results demonstrate that the adsorption constants evolved in opposite directions under the 3-h and 30-min regimes, and suggest that the differences in plaque size may be caused by variation in phage-host cell binding affinity.

### 3-h transfers select for high persistence outside the host

We mapped the mutations underpinning the small- and large-plaque phenotypes onto the ΦX174 F-protein structure. The F(T100A) substitution in the large-plaque-forming mutant evolved in the 30-min regime was located in the centre of the F monomer (**Fig. 4A**). However, the two small-plaque-associated mutations F(G321D) and F(S426*) found in isolates of the 3-h regime were at the peripheral interfaces between F monomers in the assembled capsid (**Fig. 4A)**. These peripheral mutations might influence capsid symmetry or stability by altering inter-subunit interactions [52], potentially contributing to the persistence of small-plaque-forming mutant phages during extended incubation periods, such as our 3-h regime.

**Fig. 4:**
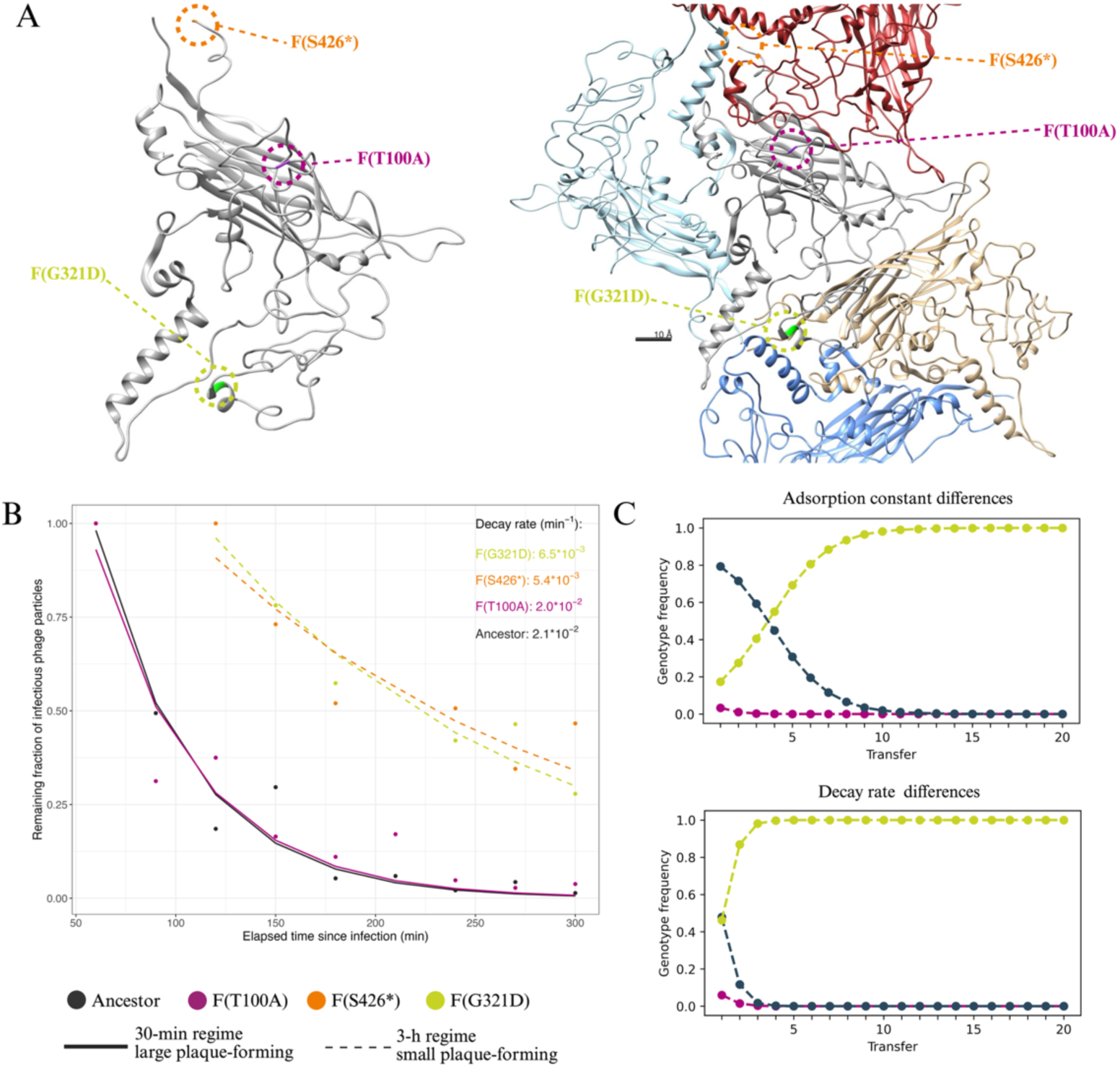
Single-point mutations in the major capsid protein modulate adsorption and persistence. (A) Location of point mutations linked to plaque size differences in the F protein monomer (left) or multimer (right) structure. Mutations F(G321D) and F(S426*) from 3-h regime isolates leading to small plaque phenotypes are shown in green and orange, respectively. The 30-min regime mutation F(T100A) leading to a large plaque phenotype is shown in purple. Mutations associated with the mall plaque phenotype are localised at monomer-monomer interfaces, whereas mutation F(T100A) is located at the F protein centre. The protein structure, resolved by McKenna *et al.* [32], was taken from RCSB PDB (reference 2BPA: https://www.wwpdb.org/pdb?id=pdb_00002bpa). The figure was prepared in UCSF Chimera (version 1.18). (B) In vitro measured decay of infectious phage particles of the ancestor (grey), large-plaque mutant T100A (purple), and small-plaque mutants G321D (green) and S426* (orange). The decay is visualised as the remaining fraction of infectious particles (y-axis) relative to the maximum phage titer, reached 60 minutes (ancestor and F(T100A)) or 120 min (F(G321D) and F(S426*)) post-infection initiation. The loss of infectious particles is shown for a five-hour infection period (x-axis). The full phage population dynamics, including the productive infection period, used to calculate the infection phage particle fraction, are shown in **Fig. S6**. See **Supplementary Methods** for model selection and parameter extraction. (C) Simulations of free phage population dynamics (y-axis) across transfers (x-axis) for the 3-h regime, incorporating differences in adsorption constants (top) or decay rates (bottom) according to values shown in Table 1 for the ancestor (grey), 30-min mutant T100A (purple), and 3-h mutant G321D (green). The simulations started with a genotype frequency of 99% of the ancestral phages and 0.5% of each mutant phage. The graphs show the predicted frequency of each type at the end of each transfer.

To test this, we performed in vitro time-course phage-bacteria co-culturing using the ancestral ΦX174, the large-plaque mutant F(T100A) from the 30-min regime, and the two small-plaque mutants (i.e. F(G321D and F(S426*)) evolved in the 3-h regime. Co-cultures were initiated at an MOI_input_ of ∼0.1, and infectious phage particles were quantified at regular intervals over five hours following chloroform extraction. All phage populations exhibited an initial productive phase, during which infectious particles increased, followed by a decay phase after the depletion of susceptible host cells (**Fig. S6**). The productive phase differed slightly between the different phages. The ancestor and the large-plaque-forming mutant evolved in the 30-min regime reached their maximum titer of infective phages ∼60 minutes post-infection, with a ∼5,000-fold increase in population size. Small-plaque mutants evolved in the 3-h regime peaked slightly later but achieved a 10,000-fold increase in population size. The higher population size peak of small-plaque mutants may be attributed to their higher adsorption constant, allowing for earlier infection initiation. The most striking differences, however, were observed during the decay phase (**Fig. 4B**). Both the ancestor and the large-plaque-forming mutant evolved in the 30-min regime showed a drastic reduction in infective phage particles, with titers declining more than 15-fold by the end of the experiment. In contrast, the small-plaque mutants evolved in the 3-h regime maintained a relatively constant number of infective particles throughout the remainder of the assay, showing only a minor decline over time.

### In silico, adsorption and persistence contribute to the selection of small-plaque mutants in the 3-h regime

To explore the roles of absorption rate and persistence in phage fitness under different transfer regimes, we developed a mathematical model of phage infection dynamics. The model captures the interaction between the bacterial host and phage with different adsorption constants or decay rates during the different transfer experiments. Upon adsorption of a phage to an uninfected host cell, the cell moves into the infected class. Phages of all types continue to adsorb to already infected cells, but only phages corresponding to the type that infected first are released upon lysis. We explicitly consider the lysis time, reflecting the time delay between adsorption of a phage and lysis of the infected cell, triggering the release of new viral particles. The full model is given in the **Methods**.

We started by exploring the role of the absorption constant in the 3-h regime. We simulated 3-h transfers with phages with an intermediate adsorption constant representing the ancestral phage, and introduced phages with higher and lower adsorption constants at low frequency to monitor changes in their frequencies throughout the simulations. All other life-history traits, including persistence, were held constant (**Table 1**). The simulations predicted an increase in the frequency of fast adsorbing phages at the expense of phages with intermediate and low adsorption constants (**Fig. 4C, top graph**), indicating that fast adsorption could explain the evolution of the small plaque-mutants under the 3-h regime.

Next, we tested if differences in decay rates could also explain the evolution of the small-plaque mutants in the 3-h regime. We incorporated the in-vitro decay rates into the in-silico 3-hour serial transfer model (**Table 1**), this time keeping adsorption constant at intermediate levels. The simulation predicted the rapid increase in the frequency of phages with higher persistence (**Fig. 4C, bottom graph**), indicating that increased persistence provides a larger fitness gain than increased adsorption in the 3-h regime. Thus, the F(G321D) and F(S426*) mutations evolved in the 3-h regime are positively pleiotropic and enhance fitness by increasing both adsorption and persistence.

### The adsorption optimum reflects a within-transfer trade-off between productive and unproductive adsorption in the 30-min regime

While the evolution of fast adsorption and increased persistence in the 3-h regime is intuitively understandable, the evolution of a lower adsorption constant under the 30-min regime is less intuitive. We expected fast adsorption to be beneficial in well-mixed liquid cultures as it would allow rapid infection of host cells [20, 21]. Hence, we were surprised that phages with dramatically reduced adsorption constants were selected under the 30-min regime.

We used our model to explore the role of the adsorption constant on fitness under the 30-min regime. First, we simulated competition between the ancestral phage and hypothetical mutants with adsorption constants ranging from the ancestor value to 1000-fold lower over a single 30-min transfer (**Fig. 5A**). Each simulation began with a 1:1 ratio of ancestral and mutant phages at an input multiplicity of infection (MOI_input_) of 0.1, and predicted the relative mutant titer after 30 minutes, corresponding the first 30-min transfer of our regime. The simulation predicted an optimal adsorption constant approximately 6-fold lower than that of the ancestor, very close to the value observed in the F(T100A) mutant. This optimum was robust across a range of burst sizes and lysis times that allow fewer than two infection cycles (**Fig. S2**). In silico, mutants with very low adsorption constants performed poorly due to limited infection initiation, whereas mutants with very high adsorption also performed poorly because rapid adsorption increases loss to binding of already infected cells, which acted as a superinfection sink. As a result, slower-adsorbing phages were predicted to remain free at a higher frequency at the end of the transfer (**Fig. 5B**) and therefore be available to initiate infections after dilution into fresh hosts. Consistent with this, in vitro measurements showed a higher number of free phages for slow-adsorbing mutants evolved in the 30-min regime than for ancestral phages (**Fig. 5C**).

**Fig. 5:**
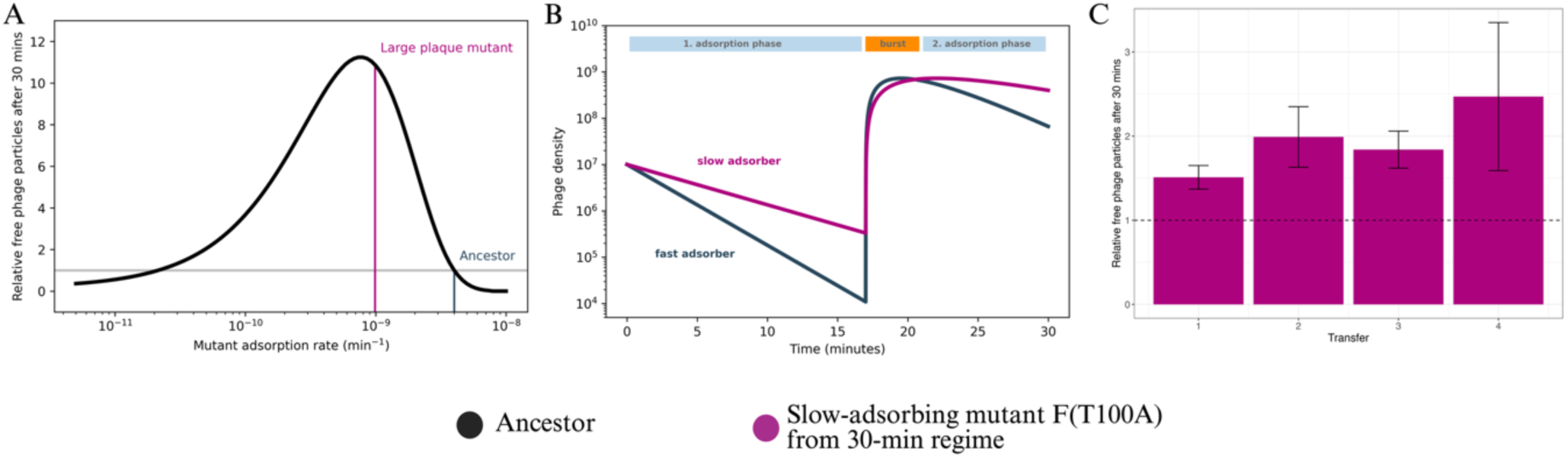
Simulations and in-vitro quantification of phage populations over single 30-min transfers. (A) Predicted optimal adsorption constants based on simulated competitions between the ancestor and hypothetical adsorption constant mutants (x-axis) over a single 30-min transfer, starting from an initial 1:1 ratio. The y-axis shows the predicted mutant-to-ancestor titer ratios after one transfer. The adsorption constant of the F(T100A) mutant evolved in the 30-min regime (purple line) is close to the predicted optimum. (B) Simulation of free phage densities (y-axis) within a single 30-min transfer (x-axis) for the ancestor (grey) and the slow-adsorbing mutant F(T100A) (purple). The free phage dynamics within one transfer predict a higher number of free phages of the slow-adsorbing mutant at the end of a transfer. The 30-min transfer is separated into a first and second adsorption period, interrupted by a burst phase, based on the in vitro determined lysis time of 17 minutes (**Fig. S3**). (C) In-vitro measured free phages of the slow-adsorbing F(T100A) mutant (purple bars) relative to ancestral free phages (y-axis) after consecutive 30-min transfers (x-axis). The number of free phages of mutant F(T100A) was normalised to the free phages of the ancestor (dashed black line). Data represent the average and standard error of three independent biological replicates. Individual titer measurements for mutant F(T100A) and the ancestral phage after each 30-min transfer are shown in **Fig. S7**.

To explore this further, we simulated phage dynamics over multiple consecutive 30-min transfers (**Fig. 6A**). Within-transfer dynamics converged after two transfers to a repeating pattern that recurred from transfer to transfer, characterized by a major burst of phage release and two phases within each transfer: an early phase with high host availability, in which many phages attach to uninfected cells and can generate progeny (hereafter, the productive phase), followed by a longer phase with low host availability, in which additional adsorption increasingly target cells that are already infected and, in our model, does not contribute to progeny production (hereafter, the unproductive phase) (**Fig. 6B-D**). The sharp separation between these phases is likely accentuated by the deterministic assumptions of our model, including a fixed 17-min lysis time. In vitro, lysis times are probably more broadly distributed, which would smooth the transition between phases. Regardless, this within-transfer structure imposes opposing selection pressures on adsorption. Faster adsorption increases productive infections during the productive phase, whereas lower adsorption reduces losses to an unproductive adsorption sink during the late unproductive phase. Hence, the optimal adsorption constant depends on the balance between these opposing selective pressures and should vary with the strength and duration of each phase under a given transfer protocol.

**Fig. 6:**
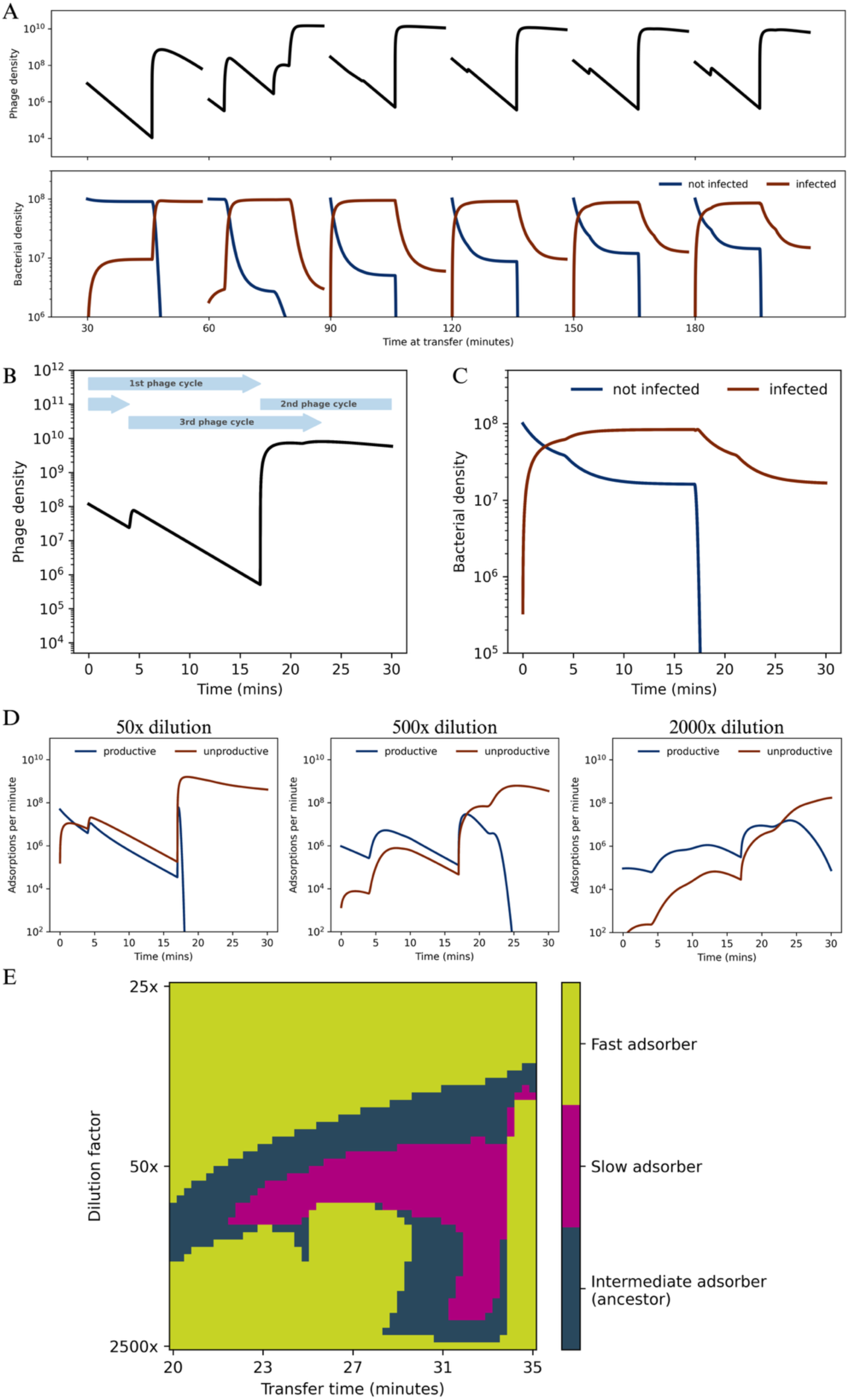
The relative strength of productive and unproductive adsorptions modulates the optimal adsorption constant in serial transfers. (A) Simulation of phage (top) and bacterial (bottom) densities (y-axis) over six 30-min transfers (x-axis) in which 2% of the bacteria-phage co-culture of the previous transfer is propagated to fresh bacteria every 30 minutes. The densities of not infected (blue) and infected (brown) bacterial cells are shown. The simulated phage and bacteria dynamics reach a stable within-transfer pattern that then repeats transfer after transfer. (B) and (C) Density (y-axis) of free phage (B) and non-infected (blue) and infected (brown) host cells over the stable 30-minute transfer pattern that emerges after a few transient transfers. The pattern is characterized by overlapping infection cycles that produce three phage burst events: a major burst after ∼17 min originating from phages that infected host cells at the beginning of the current transfer (first infection cycle), another after ∼4 min originating from the lysis of virocells infected in the previous transfer (2nd infection cycle), and another after ∼21 min originating from infections from phages that infected after the first burst (3rd infection cycle). (D) Productive (blue) and unproductive (brown) adsorptions per minute (y-axis) during a stable 30-minute co-culturing period (x-axis) for regimes with different dilution factors between transfers: 50-fold (left), 500-fold (middle), and 2000-fold (right). See also **Fig. S8**. (E) Adsorption fitness landscape across a range of transfer times (x-axis) and transfer dilution factors (y-axis). Phages with high (green), intermediate (blue) or low (purple) adsorption constants were competed in silico under transfer regimes with different co-culturing times and dilution factors between transfers. Competitions were initiated with equal frequencies of the three phages. The colour corresponding to the type fixed after 100 transfers is indicated in the plot.

We reasoned that changing the dilution factor between transfers should modify the relative weight of the early productive phase versus the late unproductive phase, thereby shifting the adsorption optimum (**Fig. 6D; Fig. S8**). To test this, we simulated competitions between phages with different adsorption constants (low, intermediate, and high) under regimes with different dilution factors and transfer times, and mapped the fitness landscape (**Fig. 6E**). As expected, different dilution factors favoured different adsorption strategies. At high dilution, fast adsorption was generally favoured because each transfer begins with abundant susceptible hosts and relatively low phage densities, increasing the payoff of quickly initiating productive infections. At intermediate dilution, where the early productive phase and the late unproductive phase are more comparable in duration and intensity, phages with intermediate and even low adsorption constants were favoured. At very low dilution, fast adsorption again became advantageous in our simulations because transfers begin with high phage densities, susceptible host cells are rapidly depleted, and there is a strong selection to capture the short initial window in which productive infections are still possible. These outcomes should be interpreted in the context of model assumptions. In particular, we do not include potential additional adsorption sinks such as cell debris, which could further penalise fast-adsorbing phages, especially at low dilution factors.

Overall, the simulations indicate that selection on adsorption can be tuned by transfer-regime parameters that alter the relative contribution of productive adsorption early in the transfer versus adsorption losses later in the transfer.

## Discussion

Bacteriophage life-history traits are commonly studied using experimental evolution under a variety of selection pressures, both in vitro and in silico [26, 27, 53–61]. Variation in host supply is a frequently used selection pressure providing limited or excess hosts at the onset of phage-bacteria co-culturing in serial-transfer experiments [2, 4, 62, 63]. In these experiments, evolution often targets the latency period and burst size, with excess hosts favouring shorter latency periods to maximise the rate of host exploitation, whereas host limitation selects for higher burst sizes to maximise yield per infection. Selection for an increased burst size often comes at the cost of longer latency periods, and vice versa, due to a lysis time-burst size trade-off [1, 64–67]. Our transfer regimes begin with hosts in excess, but host availability still changes markedly within transfers. In the 3-h regime, phages deplete susceptible hosts and must persist outside the host for much of each transfer, whereas in the 30-min regime, hosts are initially abundant but most become infected during the transfer. Our experimental results and simulations indicate that adaptation to our transfer regimes is not mediated by changes in latency period and burst size, but instead by the modulation of adsorption and persistence (**Fig. 3**; **Fig. 4**). We will dissect the reasons for this in the next paragraphs.

Longer transfer times select for phages with higher adsorption constants and greater ability to persist outside the host. This pattern can be explained by the ecological dynamics of the 3-h regime: given lysis time and host availability, multiple infection cycles can occur per transfer, leading to rapid host exhaustion. Fast-adsorbing phages initiate these cycles more efficiently. Moreover, since phage populations increase by more than two orders of magnitude per infection cycle and transfers are performed at an MOI_input_ of ∼0.1, the vast majority of sensitive host cells are lysed after two cycles, forcing phages to persist in the absence of host cells for extended periods before being supplied with new susceptible hosts (**Fig. S5**). Thus, both enhanced adsorption and environmental persistence are selected in this regime.

Two different F mutations evolved in the 3-h regime increased adsorption and dramatically increased persistence. Both mutations were located at the monomer-monomer interface in the mature capsid (**Fig. 4A**). Changes in this region can affect capsid symmetry and stability in icosahedral phages [52, 68]. The positively pleiotropic nature of these mutations makes it difficult to disentangle the relative fitness contributions of adsorption and persistence. However, the drastic increase in persistence compared to the modest elevation in the adsorption constant suggests that persistence is the primary trait under selection. This is further supported by our simulations, which show that although both increased adsorption and persistence are favoured under the 3-h regime, selection on persistence alone leads to a more rapid evolution of small-plaque mutants than selection on adsorption alone (**Fig. 4C**).

The rapid decay of the ancestral ΦX174 under our experimental conditions, with more than 90% of infectious particles lost in less than two hours (**Fig. 4B**), contrasts with the lower decay estimates previously reported [69]. Particle loss due to attachment to debris could contribute to our observed in vitro decay. However, the rapid decay was also observed when we filtered or chloroform-extracted the phages before the decay quantification (data not shown), suggesting that decay may be an intrinsic property of our ancestral phage and not exclusively related to debris attachment. Although differences between our ancestral strain and the strains used in other studies could partially account for the discrepancy in decay rates, there may also be physiologically relevant explanations. Populations of phage λ have been reported to show a biphasic decay pattern, with a rapid decay phase immediately after bursting and a slower phase thereafter [70]. If the same applies to ΦX174, it could explain the discrepancy between our results and previous decay measurements. These earlier studies may have missed the unstable fraction and primarily captured the more stable phage fraction. Further work will be needed to clarify this point.

In contrast to the 3-h regime, the shorter transfers of the 30-min regime do not impose strong selection on persistence but instead select for drastically reduced adsorption (**Fig. 3A**). The low-adsorption mutants evolved in the 30-min regime carried an F(T100A) substitution in the major capsid protein, located in an α-helix previously implicated in host binding and specificity or temperature adaptation [23, 25, 27, 39, 71, 72]. A decrease in the adsorption constant may seem counterintuitive because it reduces the rate at which infections can be initiated. However, the selection for slow adsorption can be understood as an adaptation to the specific timing of the bottlenecks in the 30-min transfer regimes. These bottlenecks favour strategies for maintaining a high density of free phages at the time of the bottleneck (transfer in our case). One such strategy is to reduce losses to adsorption during periods when most host cells are already infected. More broadly, our simulations indicate that selection on adsorption is shaped by the relative strength of two within-transfer phases imposing opposing selection pressures: a productive phase, in which uninfected cells are abundant and fast adsorption is favoured, and an unproductive sink phase, in which most cells are already infected and adsorption primarily removes phage particles via attachment to infected cells (**Fig. 6D**). Importantly, this unproductive sink may be stronger in vitro than in our model, because additional processes not explicitly included in the model—such as adsorption to cell debris or other non-productive binding interactions [50, 73, 74]—could further increase phage losses during the unproductive phase. Modulating the relative strengths of these phases (for example, by changing the dilution factor between transfers) could, in principle, allow the selection of phages with different adsorption constants (**Fig. 6E**).

Even though the complex interplay between bottlenecks and the success of adsorption rate mutants has been shown theoretically before [75], it is not usually observed in phenotype-fitness maps of bacteria-phage interactions [76]. This is because the unexpected benefit of a reduced adsorption rate in the 30-min regime requires taking into account (i) exhaustion of host cells, (ii) unproductive adsorptions on already infected cells, and (iii) frequent population bottlenecks on the order of the lysis time of the phage. A recent theoretical study showed a similar effect resulting from the need to balance productive and unproductive phases of phage growth with respect to lysis time evolution [77].

The contrasting effects of different adsorption constants across transfer regimes suggest ecological trade-offs that may drive divergent selection in natural populations. For instance, sewage water, a major source of phages, contains both a liquid phase where bacteria grow pelagically and a semi-solid phase where bacteria form biofilms [78, 79]. High adsorption may be favoured in the liquid phase to exploit transient host availability. In contrast, slow adsorption may be advantageous in biofilms or host-rich environments to avoid attachment to already infected cells [17]. Slow adsorption could also be beneficial in particle-rich environments, where phages risk irreversible binding to inert material [20, 73, 80]. Recurrent transitions between these environments may have shaped the evolutionary plasticity of adsorption in ΦX174, where single-point mutations in the F protein can lead to changes in adsorption with profound fitness consequences in different environments.

Plaque phenotypes mirrored the contrasting selective pressures in our two regimes, with small- and large-plaque-forming phages evolving under the 3-h and 30-min regimes, respectively. In theory, plaque size reduction can result from changes in the three different life history traits: (1) extended lysis time, (2) reduction in burst size, or (3) increased (or extremely low) adsorption constant [1, 22, 48, 49, 81, 82]. In our study, the differences in plaque size may reflect adsorption differences, as we found an inverse correlation between adsorption constant and plaque size (**Fig. 3**). This pattern is consistent with previous findings in phage λ, where mutations in the host recognition protein J that increased the adsorption constant led to reduced plaque size [17, 22]. Similar observations were also made in phage U136B, where fast adsorbers often formed smaller plaques [83]. A likely explanation for this is that rapid adsorption promotes local reinfection, trapping phages at the focus of infection and preventing them from reaching uninfected regions of the bacterial lawn [22, 81].

The findings presented in this manuscript have potential practical relevance. Phages with different life-history traits may be more or less suited for treating different types of infections, such as biofilm or bloodstream infections. Our results illustrate how experimental evolution can uncover key environmental trade-offs, which may inform the selection or engineering of phages for therapy. In addition, understanding how phage life-history traits evolve under different selective pressures is essential for anticipating phage evolutionary dynamics once they are administered to patients. Thus, experimental evolution serves not only as a valuable research approach but also as a practical tool to guide the development of safer, more effective, and more predictable phage-based therapies.

## Acknowledgements

We acknowledge the Sequencing Service Team from the Max Planck Institute for Evolutionary Biology in Plön for performing the Sanger Sequencing. We thank Daniel Martens and Anja Baade for their support regarding lab logistics as well as the entire Microbial Population Biology Department for their support. We also want to thank David Rogers for helping with PCR troubleshooting and Eva Lievens and Stephen T. Abedon for the insightful discussions on viral life history evolution. MR, CB, and ORM were supported by the International Max-Planck Research School for Evolutionary Biology (IMPRS EvolBio). Finally, we would like to thank our reviews and editor for their helpful comments.

## Funding

The funding for this study was received from the Max Planck Society. Work in JLG lab was also supported by ERC starting grant 853323.

## Data availability

Raw sequencing reads used to generate figures **Fig. 2** and **Fig. S4**, plaque assay images used for plaque morphology image analysis and to generate figures **Fig. 1**, **Fig. 2**, **Fig. 3B**, **Fig. S4**, **Fig. S5,** and **Fig. S11** and raw bacteria and phage counting data and measurements of plaque area and turbidity used to generate figures **Fig. 3**, **Fig. 4B**, **Fig. 5C, Fig. S3, Fig. S5, Fig. S6, Fig. S7, Fig. S9, and Fig. S10** are found in Zenodo (doi: 10.5281/zenodo.17076220).

The analysis script used for analysing and visualising the raw counting and image analysis data is in https://github.com/OReyesMatte/Reuter_2025/tree/main/Data_analysis.

Analysis information for the simulations used to generate figures **Fig. 4C**, **Fig. 5A**, **Fig. 5B**, **Fig. 6, Fig. S2, Fig. S8** is deposited in the repository found in https://github.com/OReyesMatte/Reuter_2025/tree/main/Simulations.

A list of processed plaque assay images for which plaque area and turbidity are found in https://github.com/OReyesMatte/Reuter_2025/tree/main/Plaque_analysis.

## Supplementary material

All supplementary material is found in the file “**Supplementary.pdf**”. The file contains supplementary figures **Fig. S1-S11**, supplementary tables **Table S1-S13**, and **Supplementary Methods**.

## Conflicts of interest

The authors declare no conflict of interest.

## Authors Contributions

Conceptualisation: MR, MS, JLG, FB

Methodology: MR, MS, ORM

Investigation: MR, CV, CB, JRD

Data Curation: MR, ORM

Formal analysis: MR, ORM, MS

Validation: MR, MS

Visualisation: MR, MS

Writing- Original Draft/Review & Editing: MR, JLG, MS

Supervision: JLG, FB

## Notes

### Competing Interest Statement

The authors have declared no competing interest.

### Summary of Updates

We have updated the simulations and the interpretation of some of the results. We restructured figures 4, 5 and 6, and some supplementry figures.

