## Supplementary figures, tables and methods for "Single capsid mutations modulating phage adsorption, persistence, and plaque morphology shape evolutionary trajectories in ΦX174"

**Supplementary Materials for the manuscript "Single capsid mutations modulating phage adsorption, persistence, and plaque morphology shape evolutionary trajectories in ΦX174"**

Manuela Reuter[1], Michael Sieber[1], Octavio Reyes-Matte[1], Christina Vasileiou[1], Christopher Böhmker[1], Jordan Romeyer Dherbey[2], Frederic Bertels[1,*] and Javier Lopez-Garrido[1,*]

[1]Max Planck Institute for Evolutionary Biology, Plön, Germany
[2]University of Cambridge, Cambridge, United Kingdom

**Content**

- Supplementary figures Fig. S1-S11
- Supplementary tables Table S1-S13
- Supplementary Methods

### Supplementary Figures

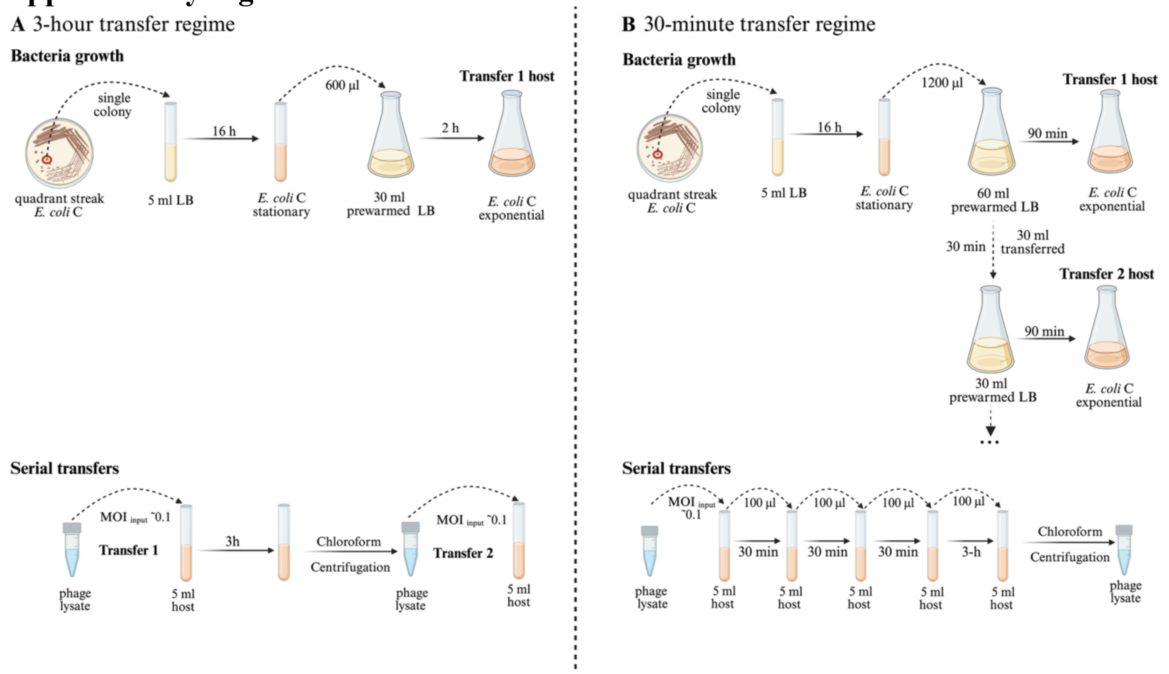


**Fig. S1: Detailed description of the 3-h and 30-min regimes.** Bacterial growth (top) and serial transfer conditions (bottom) for the (A) 3-h and (B) 30-min regime. All experimental steps were done in supplemented LB (5 mM $CaCl_2$, 10 mM $MgCl_2$). For both regimes, phage-sensitive *E. coli* C cultures were initiated from a single colony and grown for 16 h at 37°C, shaking at 250 rpm in supplemented LB. Bacterial cultures were diluted 50-fold into fresh supplemented LB and grown to mid-exponential phase. These mid-exponential cultures were used for each transfer and are hereafter referred to as "host cultures". (A) For the 3-h transfer regime, host cultures were grown for two hours before co-culturing with phage lysate at an $MOI_{input}$ of ~0.1. After 3 hours, phages were extracted using chloroform to remove host cells, and the resulting lysate was used as input for the next transfer. A single transfer was done per day, and phage concentration was determined before every transfer. (B) For the 30-minute regime, we did four 30-minute transfers followed by a single 3-h transfer per day. To maintain a continuous supply of host cultures, stationary-phase bacteria were diluted 1:50 in 60 ml of prewarmed, supplemented LB, grown for 30 min and diluted 1:2 into new flasks every 30 min to maintain exponential growth. As the 1:2 dilution eliminated the lag phase, host culture growth was shortened to 90 min to achieve a comparable physiological state growth with the 3-h regime. We generated a series of staggered cultures initiated at 30-min intervals, which were used for consecutive transfers. At each transfer, 5 ml of the appropriate host culture was aliquoted into individual tubes. For the first transfer of each day, the host culture was infected with phage lysate at an $MOI_{input}$ of ~0.1 (bottom). After 30 minutes of incubation, the phage-bacteria mix was diluted 50-fold into 5 ml of a fresh host culture. After four 30-min transfers, we performed one 3-hour co-culturing transfer and extracted the phages using chloroform. The figure was created using Biorender.com®. Reuter, M. (2025) https://Biorender.com/3c8gs9m.

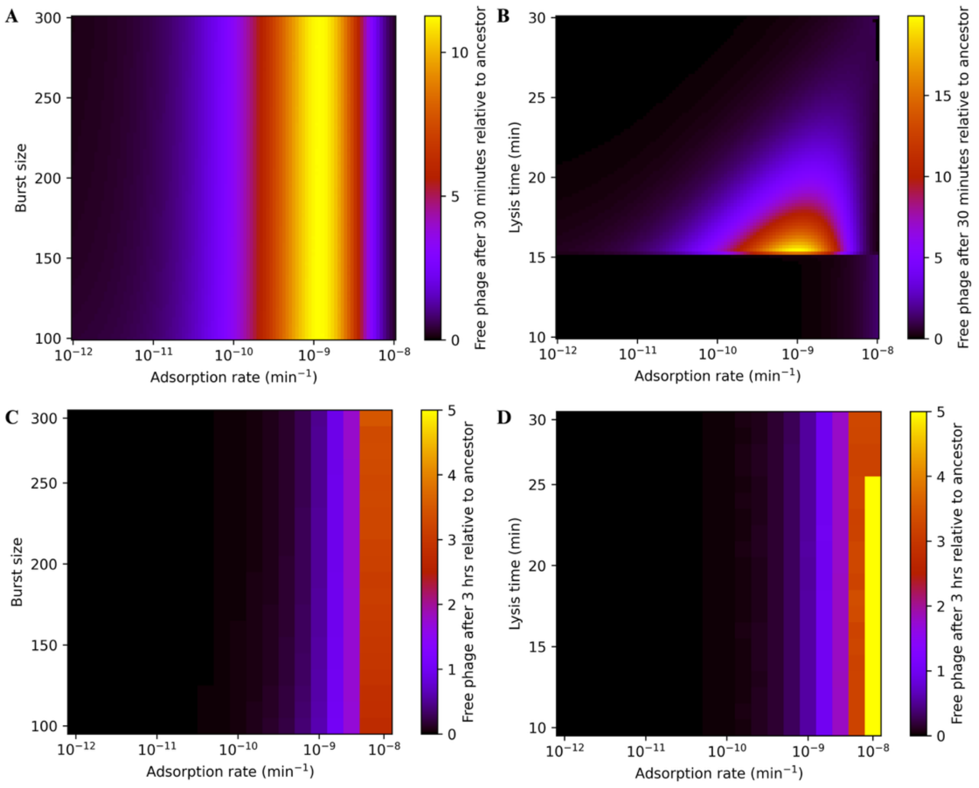


**Fig. S2: Fitness landscapes from in-silico phage competition parameter sweeps.** Graphs depict fitness landscapes of competitions between the ancestral phage and mutant phages across different adsorption rate-lysis time or adsorption-rate-burst size combinations, after a single 30-min (A and B) or 3-h (C and D) transfer. Simulations were initiated at a 1:1 mutant-to-ancestor ratio at an input MOI of 0.1, and the predicted mutant-to-ancestor ratio after 30 minutes or 3 hours was used as a proxy for fitness (indicated by the colour gradient of the heatmap), with higher values indicating higher fitness of the mutant. For all panels, adsorption rates (min$^{-1}$) are shown on the x-axis and burst size (phages released per infected cell; panels A and C) or lysis time (min; panels B and D) on the y-axis. (A) Competition simulation for a 30-min interval, across a range of adsorption rates and burst sizes. The predicted fitness peak at intermediate adsorption rates is robust across the range of burst sizes tested. For this simulation, the lysis time was kept constant (17 minutes). (B) Competition simulations for a 30-min interval across a range of adsorption rates and lysis times, with a constant burst size (200 per infected host cell), revealed two distinct areas separated at ~15 min lysis time. For lysis times longer than 15 min, which allow the completion of only one infection cycle, the simulations predict an optimal adsorption rate lower than the ancestral value and similar to that of the large-plaque mutant, with the fitness gain diminishing as lysis time increases. For lysis times shorter than 15 min, which allow the completion of two infection cycles during a 30-min transfer, higher adsorption rates become advantageous, as fast-adsorbing phages are more likely to initiate the second infection first and to produce more progeny before transfer. Given our experimentally measured lysis time of 17 min, phages with adsorption rates lower than that of the ancestor are predicted to have a fitness advantage. (C) In-silico competition simulations for a single 3-h interval, varying adsorption rate and burst size while keeping lysis time constant at 17 minutes, predict consistently high fitness for fast-adsorbing phages across all tested burst sizes. (D) Likewise, 3-h transfer competition varying lysis time also predicts high fitness for fast adsorbers across the entire range of lysis times tested.

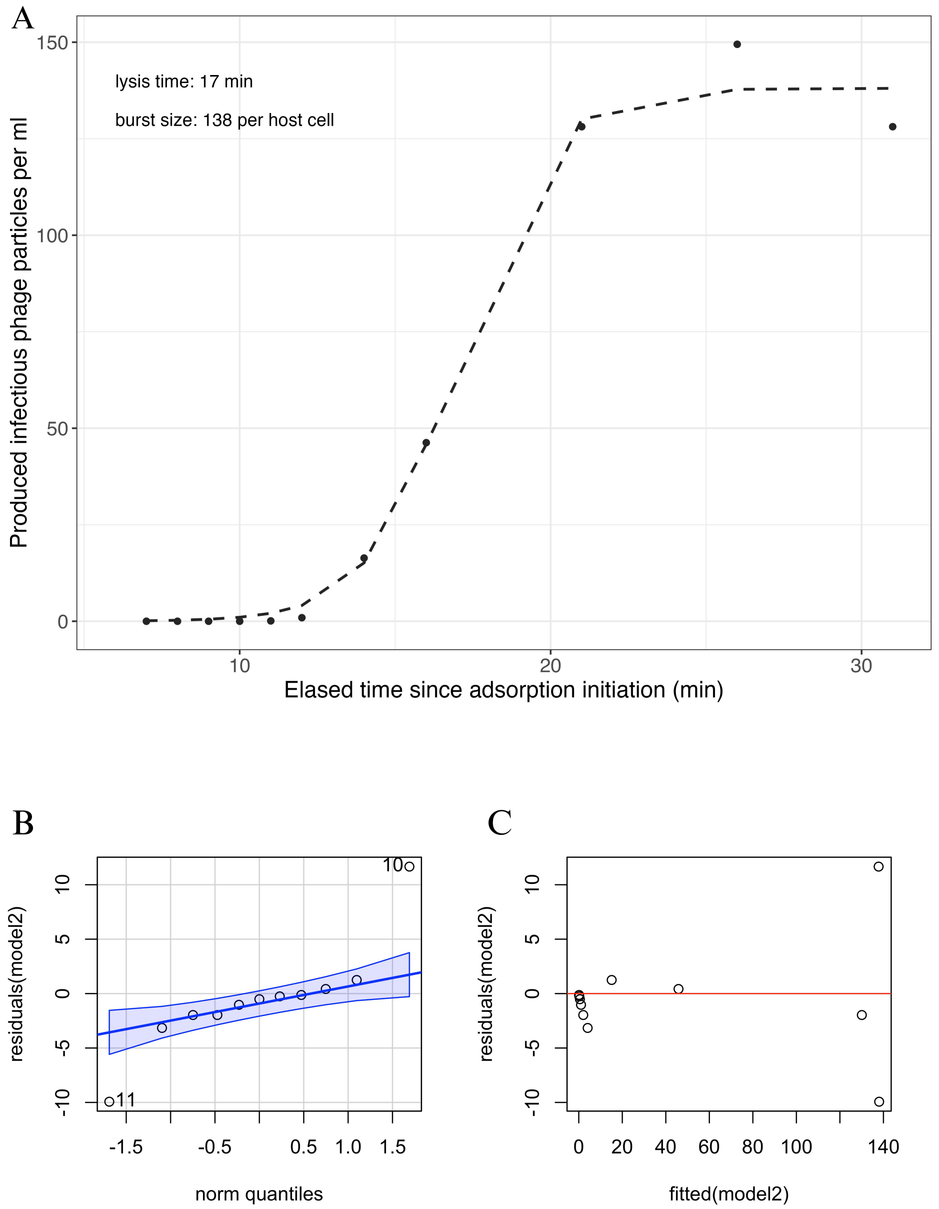


**Fig. S3: One-step growth curve to estimate lysis time and burst size of the ancestral ΦX174**. Detailed information about the one-step growth curve protocol and model fitting is provided in the **Supplementary Methods**. (A) Concentration of free phages (y-axis) over time after adsorption initiation (x-axis). Sampled timepoints are shown as black dots. The curve (dashed line) shows the predicted number of phage particles based on a logistic growth model (see Supplementary Methods for details). The parameter estimates for lysis time and burst size are shown in the top left corner of the graph (see **Tables S6 and S7** for additional details). (B) A "qqplot" was used to check for normal distribution of the logistic growth model (model2) residuals. The "norm quantiles" (x-axis) show the theoretical quantiles of a normal distribution. The residuals were normally distributed except for two outliers (numbered dots). The blue area indicates the expected uncertainty of the data distribution. (C) The "residuals vs fitted values" plot shows the fitted values of the logistic growth model (model2) on the x-axis and the model residuals on the y-axis. The values are equally distributed above and below zero (red line), and there is no correlation between residuals and fitted values.

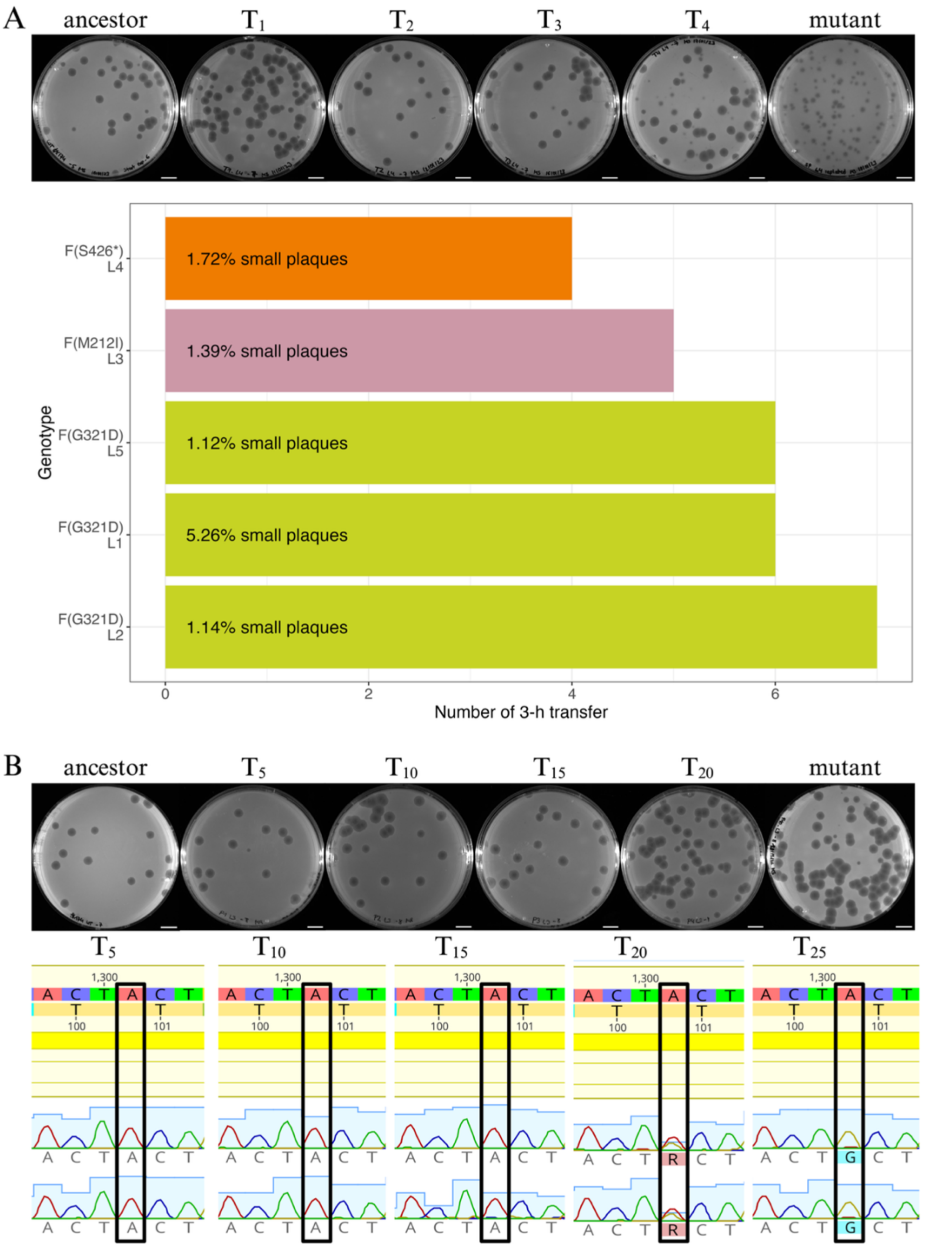


**Fig. S4: In-vitro emergence of small- and large-plaque-forming mutants in the 3-h and 30-min regime.** (A) (Top) Lysis plaques formed by one population evolving under the 3-h regime. In the 3-h regime, four transfers ($T_1$ to $T_4$) were required to observe heritable small plaques. The heritability of the plaque morphology was tested by cutting and replating individual small plaques. The last image shows the replating result of one small plaque cut from the plaque assay plate of transfer 4. Scale bars, 1 cm. (Bottom) Number of transfers (x-axis) after which small plaques started to be observed in the different 3-h evolving lines in vitro. The line number (L) and the specific genotype evolved in each case (also indicated by the colour of the bars) are shown on the y-axis. Three lines evolved small-plaque mutants carrying the mutation F(G321D) (green), one line evolved mutants carrying the substitution F(M212I) (pink), and another evolved mutant carrying the F(S426*) mutation (orange). The estimated frequency of the small-plaque mutants in the population is shown inside each bar. Frequencies were calculated based on the number of verified heritable small plaques compared to the total number of plaques on the plates. (B) (Top) Lysis plaques evolution under the 30-min regime. For the 30-minute regime, phages were plated every fifth transfer ($T_5$ to $T_{20}$). The last image of a plate full of large plaques, after forty 30-min transfers. Scale bars, 1cm. (Bottom) Sanger sequencing results of position A1301 (black box) in the phage populations from transfers in the 30-min regime. The top, yellow-shaded sequence shows the ancestral genome sequence, while the bottom half shows the chromatograms of the population sequencing. Sequencing was done in duplicates using the reverse primer ΦX1500R (**Table S1**). Transfers $T_5$-$T_{15}$ show the ancestral base "A". At transfer $T_{20}$, the single-point mutation at position A1301, giving rise to the large-plaque mutant F(T100A), occurred at a detectable frequency, seen by an additional peak indicating an "A" to "G" base change in position 1301. For transfer $T_{25}$, only the peak of base "G" is detectable, suggesting an increase in frequency of the mutation in the phage population. Note that the amino acid number shown in the figure shows the codon, not the residue number, which is shifted by -1. Sequencing figure generated Geneious version 2023.2 created by Biomatters. Available from https://www.geneious.com.

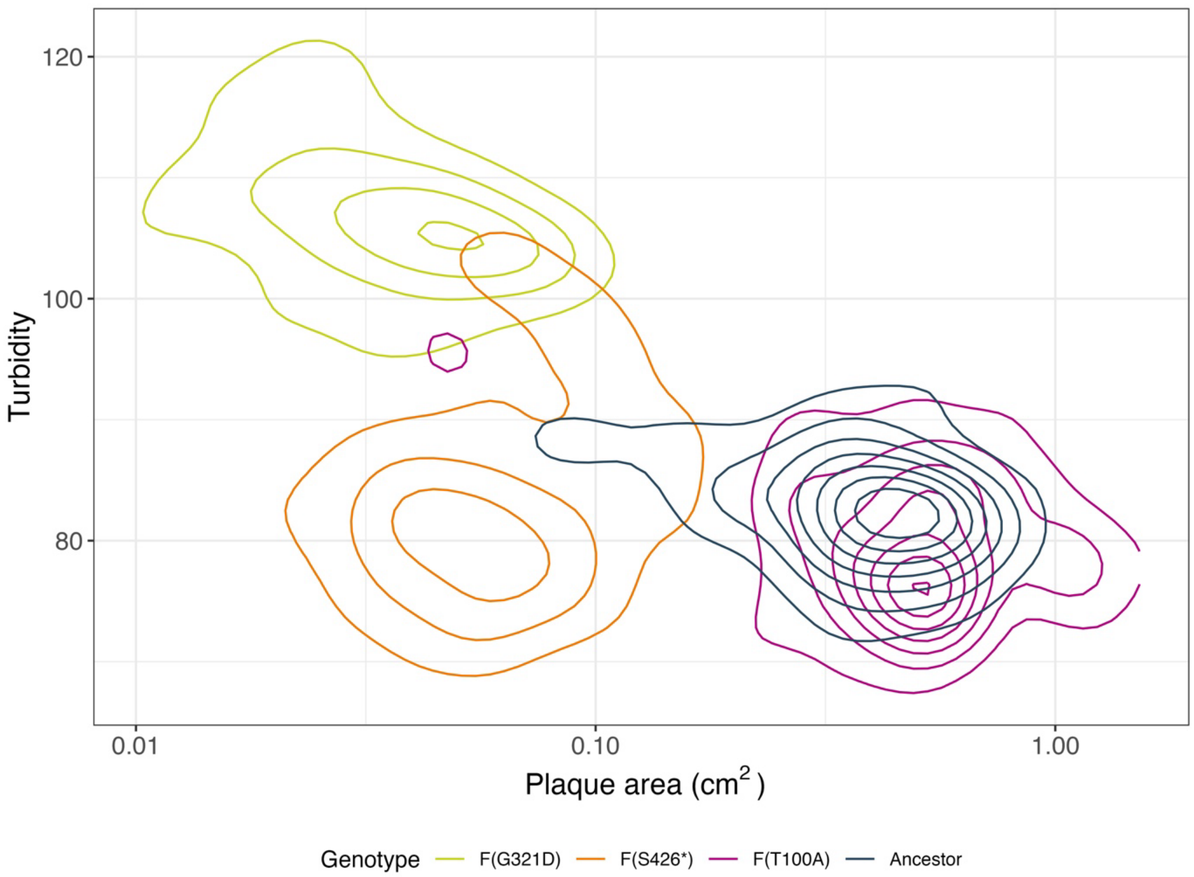


**Fig. S5: In-vitro morphological characterisation of large and small plaques.** Density plot showing the plaque area (in $cm^2$, x-axis) vs turbidity (arbitrary units, y-axis). Turbidity was estimated from plaque image intensity, with high values indicating turbid plaques and low values indicating clear plaques. Contour lines indicate clusters of plaques with size and turbidity within the range delimited by the contour. Measurements of plaque area and intensity are grouped by phage genotype: the ancestral strain (grey), the large-plaque mutant F(T100A) evolved in the 30-min regime (purple), and small-plaque mutants F(G321D) and F(S426*) isolated from the 3-h regime (green and orange, respectively). The following number of plaques was analysed for each strain: ancestor, 76 plaques; F(T100A), 60 plaques; F(G321D), 303 plaques; F(S426*), 431 plaques. The small-plaque mutants F(G321D) and F(S426*) are clearly separated from the ancestor and the F(T100A) mutant along the plaque area axis. Plaques formed by F(T100A) and the ancestor overlap substantially, although the F(T100) plaques are slightly larger and clearer. The two small-plaque further distinguished by plaque turbidity: the F(G321D) mutant forms more turbid plaques than the F(S426*) mutant. A few plaques produced by the ancestor and F(T100A) mutant overlapped in area with those of the small-plaque mutants. This likely reflects phenotypic plasticity caused by delayed infection initiation, rather than underlying genetic changes. We verified this by isolating the small plaques and replating them. In all cases, phages isolated from small plaques produced large plaques upon replating, consistent with the original genotype. Plaque area measurements were also used to generate figure **Fig. 3B**.

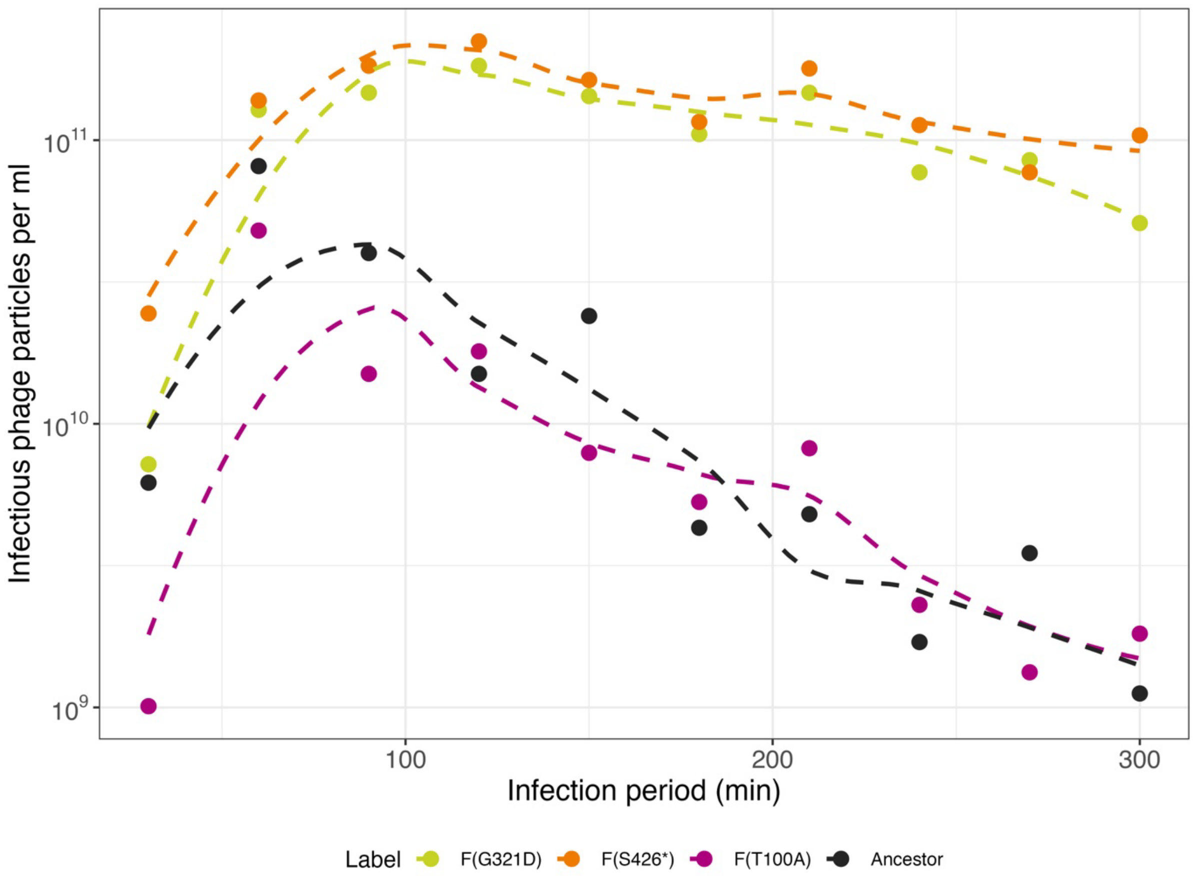


**Fig. S6: Phage population dynamics in vitro during a five-hour incubation period for different phage genotypes.** Infectious phage particles per ml (y-axis) are plotted as individual dots for each sampling time point during the 5-hour incubation period (x-axis). Each genotype is represented in a different colour: The ancestor in dark grey, the large-plaque-forming mutant F(T100A) evolved in the 30-min regime in purple, and the small-plaque-forming mutants F(G321D) and F(S426*) evolved in the 3-h regime in green and orange, respectively. The dashed lines represent fitted curves using the "loess fit" function in ggplot2 (**Table S4**). The incubation period can be divided into a growth period, in which phage titers increase due to replication in host cells, and a decay period, during which the number of phage particles decreases after susceptible host cells are exhausted. The data used to generate this figure was also used for the decay rate fitting visualised in **Fig. 4B**.

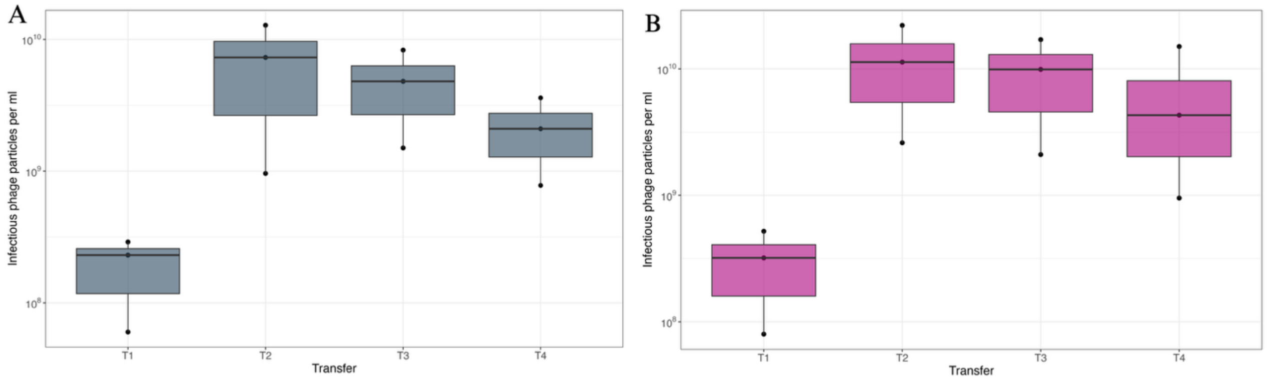


**Fig. S7: Infectious phage particle production after 30-minute transfer times for (A) ancestral ΦX174 and (B) slow-adsorbing mutant F(T100A) evolved in the 30-min regime.** For both panels, free phages were sampled and filtered before titering. Black dots show the phage titers (y-axis) after four consecutive 30-minute transfers, T1 to T4 (x-axis), for three biological replicates of the wild type (A) or the slow-adsorbing mutant F(T100A) evolved in the 30-min regime (B). Transfer T1 was initiated from phage lysate added to a final concentration of $10^7$ PFU/ml for all replicates ($MOI_{input}$~0.1). All consecutive transfers were initiated by diluting phages-bacteria co-cultures from the previous transfer by 50-fold. Boxes indicate the $25^{th}$ to $75^{th}$ percentiles, and black solid lines show the medians. Black dots represent individual biological replicates. Phage particle counts shown in this figure were also used to generate **Fig. 5C**.

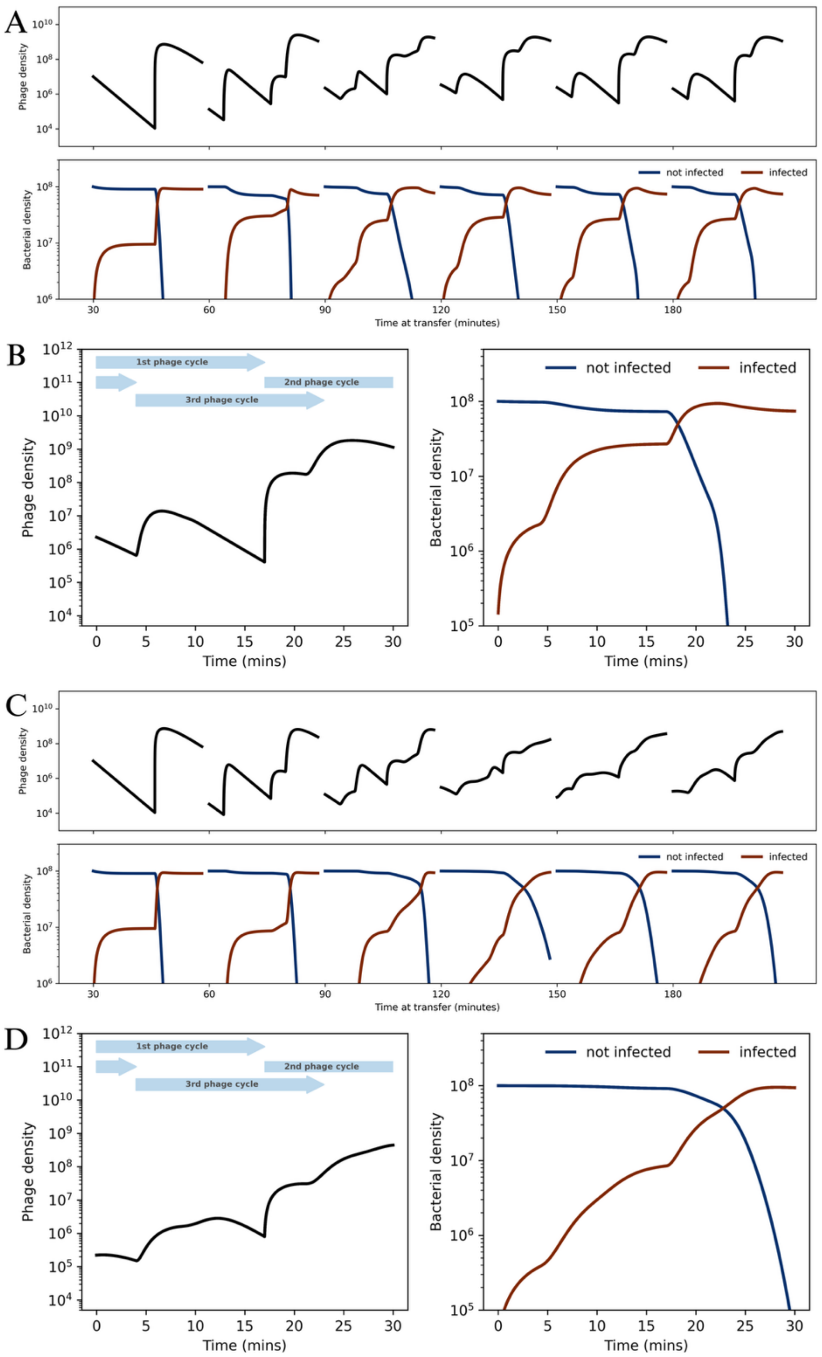


**Fig. S8: Simulations of infection dynamics during multiple 30-min transfers with different dilution factors between transfers.** (A and C) Simulation of consecutive 30-min transfers with 500-fold dilution (A) and 2000-fold dilution (C) of the bacteria-phage co-culture between transfers. The density of free phages is shown in the upper graph, and the density of not infected (blue) and infected (brown) bacterial cells is shown in the bottom graph of each panel. The series was initiated by introducing phages at an MOI of 0.1, and a stable within-transfer pattern was reached after a few transfers. (B and D) Density of free phage (left graph of each panel) and non-infected (blue) and infected (brown) host cells (right graph of each panel) over a stable 30-minute transfer in regimes with 500-fold (B) or 2000-fold (D) dilution between transfers. The patterns are characterised by overlapping infection cycles that produce three phage burst events (see legend of **Fig. 6** for details).

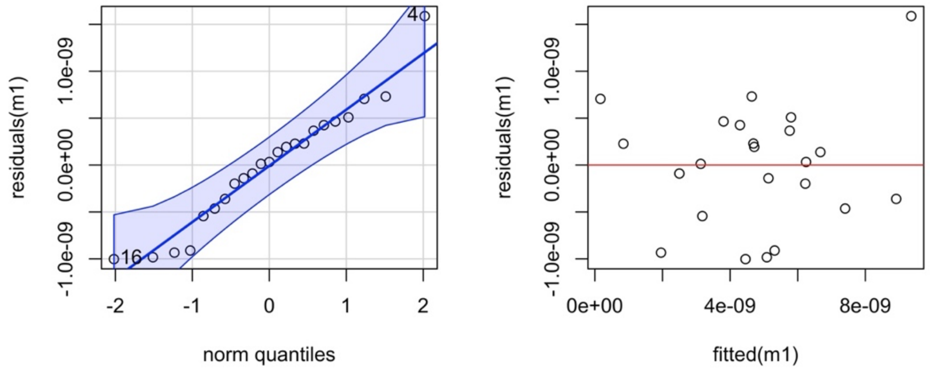


**Fig. S9: Model validation for the linear mixed effects model (m1) testing for significant differences in the adsorption constant.** (A) A “qqplot” was used to check for normal distribution of the model residuals (y-axis) compared to the theoretical quantiles of a normal distribution (x-axis). The model’s residuals were normally distributed. Two outliers were identified (numbered points), but both were still within the expected variability of the data distribution (blue shading). (B) Plot representing the residuals (y-axis) vs fitted values (x-axis) shows the fitted values of the model on the x-axis and the model residuals on the y-axis. The values are equally distributed above and below 0 (red line), and no correlation between the two variables could be found.

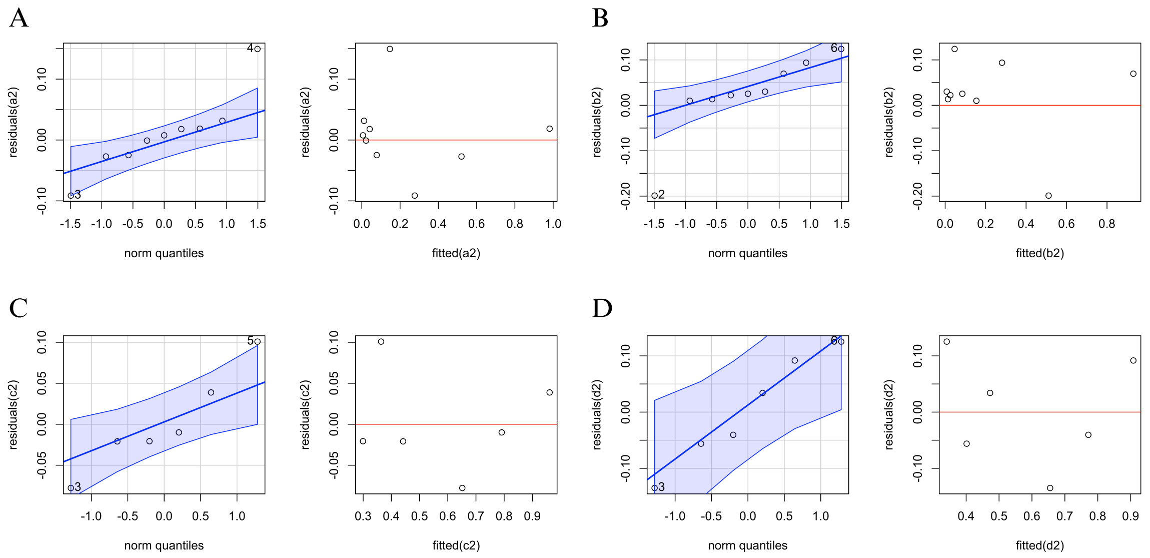


**Fig. S10: Model validation of the exponential decay fitting for the different phage genotypes.** Graphical validation of the exponential decay models (a2, b2, c2, d2) for the ancestor (A), large-plaque-forming mutant F(T100A) (B) evolved in the 30-min regime, and the small-plaque-forming mutants F(G321D) (C) and F(S426*) (D), both evolved in the 3-h regime. In each panel, a “qqplot” (left) is shown to assess the normality of the residuals (y-axis) compared to the theoretical quantiles of a normal distribution (x-axis). The blue shaded area indicates the confidence area. Outliers are indicated as numbered dots. A corresponding “fitted vs residuals” plot (right) examines the independence between fitted values and residuals. All “qqplots” show that residuals follow a normal distribution, with most datapoints falling within the confidence area. Additionally, the “residuals vs fitted” plots show no correlation between the two variables, with a similar number of points above and below the 0-line (red line) in most cases.

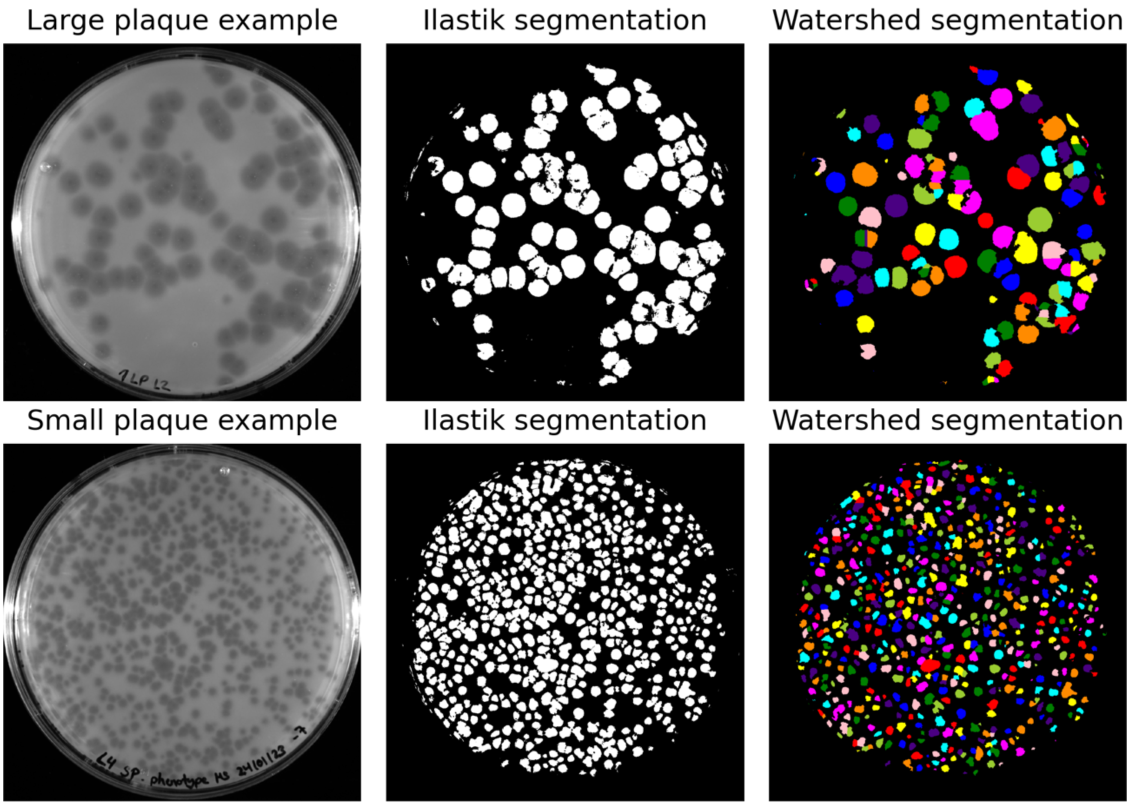


**Fig. S11: Process of getting a segmentation mask from raw plaque assay images.** The left column shows the raw plaque assay images (ChemiDoc: Bio-Rad, California, USA; 0.1s exposure time) of a large (top) and a small (bottom) phage genotype. The middle column shows the segmentation masks returned from Ilastik, showing the separation between background (black) and object of interest (white; in our case, plaques) based on the previous raw images. The right columns show corresponding segmentation masks after post-processing. The different colours indicate different plaques, which are assigned an individual ID, allowing the extraction of plaque area and turbidity measurements.

#### Supplementary Tables

**Table S1: Oligonucleotides for amplification and Sanger sequencing PCRs.** Oligonucleotides are found in the column "name", where "F" and "fw" indicate "forward" and "R" and "rev" indicate "reverse". The column "PCR" indicates if the primer was used for amplification or sequencing PCR. Tm (Q5): Melting temperature for 2x Q5® High Fidelity Master Mix (NEB, Frankfurt am Main, Germany). Ta (Q5): annealing temperature for 2x Q5® High fidelity Master Mix.

| Name | 5' - 3' Sequence | PCR | Tm (Q5) | Ta (Q5) | Reference |
|---|---|---|---|---|---|
| **ΦX0F** | GAGTTTTATCGCTTCCATG | Amplification/ Sequencing | 58°C | 59°C | [1] |
| **ΦX2953R** | CCGCCAGCAATAGCACC | Amplification/ Sequencing | 67°C | 59°C | [1] |
| **ΦX2605F** | CAGGTTGTTTCTGTTGGTGCTG | Amplification/ Sequencing | 67°C | 59°C | [1] |
| **ΦX379R** | CTTGACTCATGATTTCTTACC | Amplification/ Sequencing | 58°C | 59°C | [1] |
| **ΦX500R** | GTTACTCGTCAGAAAATCG | Sequencing | 57°C | 60°C | [1] |
| **ΦX1046R** | AGGAAGCCAAGATGGGA | Sequencing | 63°C | 60°C | [1] |
| **ΦX1500R** | TTGAGATGGCAGCAACGG | Sequencing | 66°C | 60°C | [1] |
| **ΦX2000R** | CGGAAAACATCCTTCATAGAA | Sequencing | 59°C | 60°C | [1] |
| **ΦX2500R** | TCAAACTATCAAAATATAACGTTGACGATG | Sequencing | 63°C | 60°C | [1] |
| **ΦX3381R** | GATTCTCAAATCCGGCG | Sequencing | 60°C | 60°C | [1] |
| **ΦX3849R** | CGGCAACAGCTTTATC | Sequencing | 57°C | 60°C | [1] |
| **ΦX4292R** | CACATTGTAGCATTGT | Sequencing | 52°C | 60°C | [1] |
| **ΦX5199R** | GGATTAAGCACTCCGTGGA | Sequencing | 64°C | 60°C | [1] |
| **Φx4753 correctR** | GCCTGAGTATGGTACAGC | Sequencing | 62°C | 60°C | [1] |
| **ΦX491F** | CGATTTTCTGACGAGTAAC | Sequencing | 66°C | 60°C | This study |
| **ΦX831R** | ACATCACTCCTTCTGC | Sequencing | 58°C | 60°C | This study |
| **ΦX1429F** | CCGTACCGAGGCTAACC | Sequencing | 65°C | 60°C | This study |
| **ΦX5099F** | GCTGTCGCTACTTCCCAAG | Sequencing | 66°C | 60°C | This study |
| **ΦX129R** | CCGCCAGCAGTCCACTTC | Sequencing | 69°C | 60°C | This study |
| **ΦX174_H_Sanger_fw** | GTGGCGCCATGTCTAAATTG | Sequencing | 64°C | 60°C | [2] |
| **ΦX174_F_fw** | CGCTCGTCTTTGGTATGTAGGTGG | Sequencing | 70°C | 60°C | [2] |
| **ΦX174_F_Sanger_fw** | CCTCATCGTCACGTTTATGG | Sequencing | 63°C | 60°C | [2] |
| **ΦX174_F_Sanger_fw2** | ACCGATATTGCTGGCGAC | Sequencing | 65°C | 60°C | [2] |

**Table S2: Composition of the PCR mix** used for the amplification (top) and Sanger sequencing (bottom) PCR.

| **Amplification PCR** Master Mix | |
|---|---|
| **Component** | **Volume added (µl) for 50 µl reaction** |
| 2x Q5 HF MM | 25.0 |
| Forward and Reverse Primer (10 µM) | 2.5 |
| Template (ΦX174 DNA) | 2.0 |
| HPLC water | 20.5 |

| **Sanger Sequencing PCR** Master mix | |
|---|---|
| **Component** | **Volume added (µl) for 10 µl reaction** |
| DNA (cleaned PCR product) (~100 ng) | 2.0 |
| Big Dye Terminator (BD) | 0.5 |
| 5x Big Dye Buffer (SD) | 2.0 |
| Primer (5 µM) | 1.0 |
| HPLC water | 4.5 |

**Table S3: Cycler settings used for the amplification (top) and Sanger sequencing (bottom) PCR.** The first column shows the temperature of each PCR step in °C, the second column indicates the corresponding time in seconds (sec) or minutes (min). Steps 2 to 4 were repeated for 18 cycles (amplification PCR) or 30 cycles (Sanger Sequencing PCR).

| **Amplification PCR** | | |
|---|---|---|
| **98°C** | 30 sec | |
| **98°C** | 10 sec | x 18 cycles |
| **57°C** | 20 sec | |
| **58°C** | 2 min | |
| **72°C** | 2 min | |

| **Sanger Sequencing PCR** | | |
|---|---|---|
| **96°C** | 1 min | |
| **96°C** | 10 sec | x 30 cycles |
| **56°C** | 15 sec | |
| **60°C** | 4 min | |

**Table S4: Packages and corresponding references used for data analysis in R version 4.2.1** [3].

| R Package | Reference |
|---|---|
| dplyr | [4] |
| tidyr | [5] |
| car | [6] |
| tidyverse | [7] |
| ggplot2 | [8] |
| gridExtra | [9] |
| ggthemes | [10] |
| plotrix | [11] |
| scales | [12] |
| ggpubr | [13] |
| viridis | [14] |
| stringr | [15] |
| multcomp | [16] |
| lme4 | [17] |

**Table S5: Mean and standard deviation of plaque area and adsorption constant (k), and plaque phenotype for the different genotypes quantified in vitro.** Both parameters were quantified for four different ΦX174 genotypes shown in the first column. Mutant F(T100A) was a large-plaque former isolated from the 30-min regime, and mutants F(G321D) and F(S426*) were small-plaque formers evolved in the 3-h regime. Mean values are the mean and the standard error (SE).

| Genotype | Plaque Size ($cm^2$) ± SE | Adsorption constant ($ml^{-1}$ $min^{-1}$) ± SE | Plaque phenotype |
|---|---|---|---|
| Ancestor | 0.375<br>± 0.021 | $4.15*10^{-9}$<br>± $5.98*10^{-10}$ | Ancestral<br>Large |
| A1301G/<br>F(T100A) | 0.518<br>± 0.039 | $9.87*10^{-10}$<br>± $6.04*10^{-11}$ | Large |
| G1965A/<br>F(G321D) | 0.033<br>±0.001 | $6.10*10^{-9}$<br>± $9.29*10^{-10}$ | Small |
| C2280A/<br>F(S426*) | 0.066<br>± 0.002 | $5.93*10^{-9}$<br>± $5.97*10^{-10}$ | Small |

**Table S6: Degrees of freedom (DF), Akaike Information Criterion (AIC), and Bayesian Information Criterion (BIC) values for the exponential and logistic growth model of the one-step growth curve fitting**. Models were based on **Equations S1.1** (exponential) and **S1.2** (logistic) (see **Supplementary Methods**). The best-fit model was selected based on the lowest AIC and BIC values.

| Model | DF | AIC | BIC |
|---|---|---|---|
| Exponential growth | 3 | 117.0 | 118.2 |
| Logistic growth | 4 | 73.8 | 75.4 |

**Table S7: Burst size and lysis time estimates derived from fitting a logistic growth model to the one-step growth curve data of ancestral phage ΦX174**. For both parameters, the estimate, standard error, t- and adjusted p-value are shown.

| Parameter | Estimate | Standard error | t-value | Adjusted p-value |
|---|---|---|---|---|
| Lysis time "l" | 17.0 | 0.3 | 48.8 | < 0.001 |
| Burst size "b" | 138.1 | 3.8 | 35.8 | < 0.001 |

**Table S8**: **Estimation of adsorption constants "k" for the different genotypes quantified in vitro.** Column "Genotype" shows the four phage genotypes (i.e. ancestral, 30-min regime large-plaque mutant F(T100A), and 3-h regime small plaque mutants F(G321D) and F(S426*), for which the adsorption dynamics were tested. The column "Experimental run" indicates the number of biological replicates done for each genotype. Slope estimate, standard error, and multiple $R^2$ values were extracted from the linear model fit for each genotype and experimental run. The adsorption constant "k" (in $ml^{-1}$ $min^{-1}$) was calculated based on **Equation S2.1** (see **Supplementary Methods**) for each genotype and experimental run separately.

| Genotype | Experimental run | Slope estimate | Standard error | $R^2$-value | Adsorption constant "k" |
|---|---|---|---|---|---|
| **Ancestor** | 1 | -0.9914 | 0.0328 | 0.999 | $4.91*10^{-09}$ |
| | 2 | -0.8397 | 0.0154 | 1.000 | $6.94*10^{-09}$ |
| | 3 | -0.7889 | 0.0318 | 0.998 | $3.14*10^{-09}$ |
| | 4 | -0.9439 | 0.2299 | 0.944 | $4.27*10^{-09}$ |
| | 5 | -0.7034 | 0.2483 | 0.889 | $2.63*10^{-09}$ |
| | 6 | -0.9422 | 0.1843 | 0.963 | $2.41*10^{-09}$ |
| | 7 | -0.9246 | 0.2358 | 0.939 | $4.72*10^{-09}$ |
| **F(T100A)** | 5 | -0.2858 | 0.0393 | 0.981 | $1.07*10^{-09}$ |
| | 6 | -0.3401 | 0.0286 | 0.993 | $8.70*10^{-10}$ |
| | 7 | -0.2003 | 0.0226 | 0.988 | $1.02*10^{-09}$ |
| **F(S426*)** | 1 | -1.2174 | 0.3038 | 0.941 | $6.03*10^{-09}$ |
| | 2 | -1.0349 | 0.0393 | 0.999 | $8.55*10^{-09}$ |
| | 3 | -1.3484 | 0.1404 | 0.989 | $5.37*10^{-09}$ |
| | 4 | -0.9737 | 0.1340 | 0.981 | $4.41*10^{-09}$ |
| | 5 | -1.3133 | 0.2973 | 0.951 | $4.92*10^{-09}$ |
| | 7 | -1.2369 | 0.3081 | 0.942 | $6.31*10^{-09}$ |
| **F(G321D)** | 1 | -1.3753 | 0.1598 | 0.987 | $6.81*10^{-09}$ |
| | 2 | -1.3246 | 0.3041 | 0.950 | $1.09*10^{-08}$ |
| | 3 | -1.0299 | 0.0057 | 1.000 | $4.10*10^{-09}$ |
| | 4 | -1.3544 | 0.4811 | 0.888 | $6.13*10^{-09}$ |
| | 5 | -1.3334 | 0.1496 | 0.988 | $4.99*10^{-09}$ |
| | 6 | -1.3502 | 0.2778 | 0.959 | $3.45*10^{-09}$ |
| | 7 | -1.2309 | 0.1115 | 0.992 | $6.28*10^{-09}$ |

**Table S9: Results of the ANOVA comparing the null (no fixed effect) with the full (fixed and random effect) model to test if the phage genotype significantly contributed to the variation in the in vitro measured adsorption constant.** Npar shows the within degrees of Freedom, AIC, and BIC, the Akaike Information Criterion and the Bayesian Information Criterion, respectively. Thε $\chi^2$-value, the degrees of freedom between groups, and the p-value are shown. The significant p-value suggests that the phage genotype significantly contributes to explaining the adsorption constant variance. Hence, we continued with the full model for further statistical analysis.

| Model | npar | AIC | BIC | logLik | deviance | $\chi^2$ | DF | p-value |
|---|---|---|---|---|---|---|---|---|
| **Null model** | 3 | -847.2 | -843.80 | 426.60 | -853.21 | - | - | - |
| **Full model** | 6 | -870.97 | -864.16 | 441.49 | -882.97 | 29.764 | 3 | <0.0001 |

**Table S10: Parameter estimates extracted from the full model for the random effect "Experimental run" (top) and the fixed effect "Genotype" (bottom).** For the random effect, the groups, name, variance and standard deviation are shown. For the fixed effects, the estimates for the four different phage genotypes are shown. Intercept is small-plaque-former F(G321D) from the 3-h regime, followed by small plaque mutant F(S426*) also from the 3-h regime, the large plaque mutant F(T100A) evolved in the 30-min regime and the ancestor. In addition to the parameter estimate, the standard error (Std. Error) and t-value are shown.

| **Random effects** | | | |
|---|---|---|---|
| **Groups** | **Name** | **Variance** | **Std. Dev** |
| **Run** | Intercept | $2.822*10^{-18}$ | $1.68*10^{-9}$ |
| **Residual** | | $6.623*10^{-19}$ | $8.14*10^{-10}$ |
| **Number of obs: 23, groups: Run, 7** | | | |
| **Fixed effects** | | | |
| | **Estimate** | **Std. Error** | **t-value** |
| **Intercept F(G321D)** | $6.102*10^{-9}$ | $7.055*10^{-10}$ | 8.649 |
| **F(S426*)** | $-4.454*10^{-10}$ | $4.586*10^{-10}$ | -0.971 |
| **F(T100A)** | $-4.292*10^{-9}$ | $5.983*10^{-10}$ | -7.173 |
| **Ancestor** | $-1.956*10^{-9}$ | $4.350*10^{-10}$ | -4.496 |

**Table S11: Results of the Tukey's HSD post-hoc test for adsorption constant differences between groups.** The columns show the pairwise comparison between two phage genotypes, parameter estimates, standard error (Std. Error), Z- and adjusted p-value. Significant p-values are indicated with an asterisk. The "Holm-Bonferroni method" was used to adjust for multiple testing.

| Pairwise comparison | Estimate | Std. Error | Z-value | p-value |
|---|---|---|---|---|
| **Ancestor-F(T100A)** | $2.336*10^{-9}$ | $5.983*10^{-10}$ | 3.904 | <0.001* |
| **F(S426*)-F(T100A)** | $3.846 *10^{-9}$ | $6.296*10^{-10}$ | 6.109 | <0.001* |
| **F(G321D)-F(T100A)** | $4.292*10^{-9}$ | $5.983*10^{-10}$ | 7.173 | <0.001* |
| **F(S426*)-Ancestor** | $1.511*10^{-9}$ | $4.586*10^{-10}$ | 3.294 | 0.002* |
| **F(G321D)-Ancestor** | $1.956*10^{-9}$ | $4.350 *10^{-10}$ | 4.496 | <0.001* |
| **F(G321D)-F(S426*)** | $4.454*10^{-10}$ | $4.586*10^{-10}$ | 0.971 | 0.331 |

**Table S12: Comparison of a linear versus an exponential decay model based on AIC (Akaike Information Criterion) and BIC (Bayesian Information Criterion).** The linear model was based on **Equation S3.1**, the exponential decay model on **Equation S3.2** (see **Supplementary Methods**). Model selection was done for each phage genotype (i.e. ancestor, large-plaque-forming F(T100A) evolved in the 30-min regime, and small-plaque-forming mutants F(G321D) and F(S426*) evolved in the 30-min regime) individually. Based on the lower AIC and BIC values, the exponential decay model was selected for model fitting for all four genotypes.

| Phage genotype | Linear model | | Exponential decay | |
|---|---|---|---|---|
| | **AIC** | **BIC** | **AIC** | **BIC** |
| **Ancestor** | 0.34 | 0.93 | -18.73 | -18.14 |
| **F(T100A)** | 0.09 | 0.68 | -12.04 | -11.45 |
| **F(G321D)** | -7.79 | -8.41 | -11.68 | -12.23 |
| **F(S426*)** | -3.55 | -4.17 | -5.90 | -6.53 |

**Table S13: Parameter estimates extracted from the exponential decay model fitted to the in vitro measured decay curve of each phage genotype.** The table includes the estimated intercept (a) and slope (b), along with their standard errors (Std. Error), t-values and p-values. Decay rates were determined for the ancestor, the large-plaque mutant F(T100A) of the 30-min regime, and small-plaque mutants F(G321D) and F(S426*) from the 3-h regime.

| Phage Genotype | Parameter | Estimate | Std. error | t-value | p-value |
|---|---|---|---|---|---|
| **Ancestor** | Intercept (b) | 3.4833 | 0.7084 | 4.917 | 0.002 |
| | Decay rate (a) | -0.0211 | 0.0027 | -7.797 | <0.001 |
| **F(T100A)** | Intercept (b) | 3.0787 | 0.9048 | 3.403 | 0.011 |
| | Decay rate (a) | -0.0199 | 0.0038 | -5.206 | 0.001 |
| **F(G321D)** | Intercept (b) | 2.0918 | 0.3087 | 6.777 | 0.002 |
| | Decay rate (a) | -0.0065 | 0.0009 | -7.518 | 0.002 |
| **F(S426*)** | Intercept (b) | 1.7450 | 0.4072 | 4.286 | 0.013 |
| | Decay rate (a) | -0.0054 | 0.0013 | -4.146 | 0.014 |

**Supplementary Methods**

In this section, we provide additional information about the methods used in this paper.

**One-step growth curve to estimate the lysis time and burst size of the ancestral ΦX174**

Lysis time (the time at which 50% of the phage population completed one full infection cycle, from adsorption to host cell lysis) and burst size (number of phage progeny produced per infected host cell) were determined for the ancestral ΦX174 using one-step growth curves. Our protocol was a modified version of a standard one-step protocol [18, 19], tailored to our phage model. The ancestral ΦX174 phage lysate was prepared as described in the Methods section, except that free phages were harvested 1 hour post-infection rather than 3 hours, to maximise phage concentration and to minimise potential phage stability problems observed in the persistence assays.

To perform the assay, a phage-sensitive *E. coli* C culture was initiated by inoculating 60 ml of pre-warmed, supplemented LB with 1.2 ml of overnight culture and incubating for 2 hours under standard conditions. After incubation, colony-forming units were quantified as described in "Phage lysate preparation and titration" in the Methods section, and phage adsorption was carried out for 6 min as described in the "Adsorption assay" but omitting the sampling steps. To prevent secondary adsorptions, the culture was diluted 2000-fold in 20 ml of prewarmed, non-supplemented LB in a 50 ml Falcon tube (114x28 mm, PP, Sarstedt AG Co. KG, Nümbrecht, Germany). Samples of 300 µl (100 µl at later time points with higher titers) were collected at specific time points until 30 minutes post-adsorption. Samples were immediately placed on ice, and infectious phages were isolated using chloroform treatment to stop the infection process and to avoid potential decay effects. Infectious phage particles were quantified by plaque assay.

To estimate burst size and lysis time, we first corrected the phage titers for unabsorbed phages that did not contribute to burst size. Then, we calculated the ratio between phage titers at the sampled time points and the susceptible host cell concentration. We fitted a non-linear model to the one-step growth curve in R 3.6.3 [3] using the "nonlinear least squares" function (nls).

To select the best model, we compared the AIC (Akaike Information Criterion) and BIC (Bayesian Information Criterion) between two non-linear models, either with exponential

(**Equation S1.1**) or logistic growth (**Equation S1.2**). For both models, "y" was the number of produced phage particles as the response variable, and "x" was the sampling time point in minutes as the fixed effect. For the exponential growth model (**Equation S1.1**), parameter "a" captured the slope and "b" the vertical deflection of the exponential growth curve. In the logistic growth model (**Equation S1.2**), parameter "b" captured the carrying capacity, used as a burst size and "l" the inflexion point of the curve as the lysis time estimate (the time point at which 50% of the phage population has been released from lysed bacteria, assuming lysis time following a normal distribution [18]. We used the "SSlogis" function to fit the logistic growth model.

$$y = b + e^{ax} \qquad \textit{Equation S}1.1$$

$$y = \frac{b}{1 + e^{(l-x)}} \qquad \textit{Equation S}1.2$$

Due to the lower AIC and BIC values, we selected the logistic growth model to fit the one-step growth curve data (**Table S6**). Parameter estimates for burst size and lysis time are found in **Table S7**. The logistic model was visualised by plotting the predicted values of the model onto the experimentally acquired data (**Fig. S3A**)

For model validation, we used a "qqplot" and a "residuals vs fitted" value plot. The "qqplot" showed normal distribution of the models' residuals (**Fig. S3B**). The "residuals vs fitted" values plot showed no observable pattern between the two plotted variables (**Fig. S3C**). Hence, our logistic growth model passed the model validation.

**Adsorption constant quantification for ΦX174 genotypes with different lysis plaque phenotypes**

The loss of free phage particles due to irreversible adsorption to host cells was quantified as described in the Methods section, a modified version of the Hyman and Abedon protocol [19] for at least three biological replicates per phage genotype: ancestor, 30-min mutant F(T100A), and 3-h mutants F(G321D) and F(S426*). To calculate the adsorption constant "k" for each genotype, we determined the slope of the adsorption curve by fitting a linear model to the $log_{10}$-transformed phage titers, capturing the adsorption dynamics over time. The model used the phage titer (plaque-forming units per ml) as the response variable, and the sampling time point (min) as the fixed effect. Model fitting was performed independently for each phage genotype and experimental run to extract replicate estimates of the adsorption slopes (**Table S8**). We used the coefficient of determination $R^2$ to capture the difference between observed and predicted values, with extracted values ranging between 0.89 and 1.00 (**Table S8**). Hence, all experimental runs were used for further analysis.

The adsorption constant "k" was calculated for each biological replicate using **Equation S2.1**, where "B" is the bacterial concentration in colony-forming units per ml (**Table S8**). The adsorption constant was extracted from the slope of every regression (a negative value), multiplied by -1 to get a positive value and divided by the bacterial concentration to normalise for differences in host cells between experimental runs.

$$adsorption\ constant\ k\ (ml^{-1}min^{-1}) = \frac{-slope}{B} \qquad Equation\ S2.1$$

To assess statistical differences in k between genotypes, we first tested the distribution of the adsorption constants using the "Shapiro-Wilk Test", which indicated normality (W= 0.9632, p=0.5309). Homogeneity of variances was confirmed using the "Levene-Test" ($DF_{group}$=3, $DF_{samples-group}$=19, F = 1.2369, p=0.324). Adsorption constant differences between phage genotypes were analysed using a linear mixed-effects model (LMM), with adsorption constant as the response variable, genotype as a fixed effect, and experimental run as a random effect to account for experimental variability. ANOVA comparing the fill model to a Null-Model (removing genotype as a fixed effect) revealed that the genotype significantly contributed to the observed variation in adsorption constants (**Table S9**). Parameter estimates for the fixed and random effects are summarised in **Table S10.**

We validated our full model graphically using a "qqplot" and a "residuals vs fitted" value plot (**Fig. S9**). The qqplot showed normal distribution of the models' residuals, and no pattern was found between residuals and fitted values of the model. Hence, the model passed the graphical validation.

Post hoc comparisons using Tukey's HSD test (**Table S11**) revealed that both small-plaque-forming mutants had a significantly higher adsorption constant than the ancestor [F(G321D): Z = 4.496, p<0.001, F(S426*: F= 3.294, p<0.002)] and the large-plaque-forming mutant F(T100A) [F(G321D): F=7.173, p<0.001, F(S426*): F= 6.109, p<0.001)]. In contrast, the adsorption constant of the large-plaque-forming mutant F(T100A) was significantly lower than that of the ancestral (F=-3.904, p<0.001).

**Quantification of infectious particle decay of ΦX174 during long co-culturing times**

We quantified the loss of infectious phage particles during a culturing period of five hours to determine phage decay dynamics (**Fig. S6**) of ancestral phage ΦX174 as well as the 30-min mutant F(T100A), and the 3-h mutants F(G321D) and F(S426*). To quantify the decay rates of individual phage genotypes, we first calculated the ratio between phage titers during the decay phase compared to the peak productivity [120 min post-infection for F(G321D) and F(S426*) and 60 min post-infection for F(T100A) and the ancestor].

To determine the decay rate of infectious particles for each phage genotype, we compared a linear (**Equation S3.1**) and an exponential decay (**Equation S3.2**) model fitted to the ratios for each of the four phage genotypes separately. The ratio was used as the response variable "y", and the sampling time point as a fixed effect "x" for both models. For the linear model, "b" captured the intercept and "a" the slope of the curve. For the exponential decay, parameter "b" captured the vertical inflexion and "a" the slope of the curve.

$$y = b - ax \qquad \qquad Equation\ S3.1$$

$$y = b + e^{-ax} \qquad \qquad Equation\ S3.2$$

The exponential decay model was selected for model fitting based on the lower AIC and BIC values for all four genotypes (**Table S12**). The parameter estimate for "a" was extracted as an estimate for the decay rate (**Table S13**). We performed a graphical validation of our model fitting using a "qqplot" checking for the normal distribution of the models' residuals and a “residuals vs fitted values” plot to check for independence between the models’ residuals and predicted “y” values. The graphical model validation was passed for all four genotypes (**Fig. S10**).

**Image analysis**

The lysis plaque image processing pipeline consists of three steps: semantic segmentation of plaques, instance segmentation of the binary mask, and feature extraction of the plaques.

Segmentation of plaques was done via training a random forest pixel classifier using Ilastik [20]. Following the available online documentation, we included all base parameters for intensity, edge, and texture features. Ten images containing different plaque morphologies were sparsely annotated, and then the live pixel classification was used to fully segment the images into two categories: “background” and “plaques”. The “background” class included agar without lysis plaques, plate edge, dark background, and space in between plaques to facilitate downstream instancing. For the “plaque” class, annotation was done on multiple small and large plaques, with the whole plaque being annotated. The classifier was then run on the images analysed for this work, getting a “simple segmentation” output which gave mask images with two values: 1 for background and 2 for plaques.

To generate instances, we performed Watershed segmentation of the binarised segmentation masks. Masks were first eroded (to remove “loose” pixels) and holes inside objects filled, then processed with the Watershed segmentation to obtain individual instances (**Fig. S11**). Next, for each image, segmentations were manually curated, where only the plaques that were sure to be single colonies were kept.

Finally, we extracted three parameters from the processed segmentations: the plaque area (converted to $cm^2$), the distance transform, and the grayscale intensity of the original plaque image by overlaying both. Next, for each plaque, we averaged the values of intensity for each unique distance transform value, giving an estimate of how intensity (proxied to turbidity) changes based on the distance to the plaque periphery.
